# Loss of function mutation of mouse *Snap29* on a mixed genetic background phenocopy abnormalities found in CEDNIK and 22q11.2 Deletion Syndrome patients

**DOI:** 10.1101/559088

**Authors:** Vafa Keser, Jean-François Boisclair Lachance, Sabrina Shameen Alam, Youngshin Lim, Eleonora Scarlata, Apinder Kaur, Zhang Tian Fang, Shasha Lv, Pierre Lachapelle, Cristian O’Flaherty, Jefferey A. Golden, Loydie A. Jerome-Majewska

## Abstract

Synaptosomal-associated protein 29 (*SNAP29*) is a member of the SNARE family of proteins involved in maintenance of various intracellular protein trafficking pathways. *SNAP29* maps to the 22q11.2 region and is deleted in 90% of patients with 22q11.2 deletion syndrome (22q11.2DS). However, the contribution of hemizygosity of *SNAP29* to developmental abnormalities in 22q11.2DS remains to be determined. Mutations in *SNAP29* are responsible for the developmental syndrome called CEDNIK (cerebral dysgenesis, neuropathy, ichthyosis, and keratoderma). On an inbred C57Bl/6J genetic background, only the ichthyotic skin defect associated with CEDNIK was reported. In this study, we show that loss of function mutation of *Snap29* on a mixed genetic background not only models skin abnormalities found in CEDNIK, but also phenocopy ophthalmological, neurological, and motor defects found in these patients and a subset of 22q11.2DS patients. Thus, our findings indicate that mouse models of human syndromes should be analyzed on a mixed genetic background. Our work also reveals an unanticipated requirement for *Snap29* in male fertility, and support contribution of hemizygosity for *SNAP29* to the phenotypic spectrum of abnormalities found in 22q11.2DS patients.

## Introduction

CEDNIK (cerebral dysgenesis, neuropathy, ichthyosis, and keratoderma) patients have mutations in *SNAP29* and present with a number of clinical manifestations, including skin defects at birth or in the first few months of life, failure to thrive, cerebral malformations, developmental delay, severe mental retardation, roving eye movements during infancy, trunk hypotonia, poor head control, craniofacial dysmorphisms, and mild deafness (Fuchs-Telem et al., 2011, Hsu et al., 2017, Sprecher et al., 2005). However, *SNAP29* mutations in human show variable expressivity and incomplete penetrance. For example, patients carrying the c.DNA486_487insA mutation can present with the full constellation of CEDNIK-associated defects (Fuchs-Telem et al., 2011) or only present with motor and neurological abnormalities: polymicrogyria, trunk hypotonia, dysplastic or absent corpus callosum, seizures, and hypoplastic optic nerves, but no dermatological abnormalities or dysmorphic features (Diggle et al., 2017, Fuchs-Telem et al., 2011). Moreover, one patient with a truncating heterozygous mutation in *SNAP29* was reported to show autosomal dominant nocturnal frontal lobe epilepsy (ADNFLE; (Sun et al., 2016)), suggesting that heterozygous mutation in *SNAP29* is associated with incomplete penetrance.

*SNAP29* maps to human chromosome 22q11.2 and is deleted in 90% of patients carrying the common 3 Mb deletion responsible for the autosomal dominant 22q11.2 deletion syndrome (22q11.2DS). We previously showed that hemizygous deletion of 22q11.2 uncovers recessive deleterious variants in the intact *SNAP29* allele resulting in ichthyosis and neurological manifestations in a subset of patients (McDonald-McGinn et al., 2013). Intriguingly two patients with mutations in *SNAP29* and hemizygous for 22q11.2 were atypical and did not have any skin abnormalities, a hallmark of CEDNIK, while only one patient had neurological abnormalities (McDonald-McGinn et al., 2013), consistent with the hypothesis that mutation in *SNAP29* results in variable expressivity and incomplete penetrance.

We postulated that a mouse model with loss of function mutation in *Snap29* on a mixed genetic background would model abnormalities found in patients with CEDNIK and 22q11.2DS, and could be used to uncover the etiology of these abnormalities. In this study, we show that constitutive loss of function mutation in *Snap29* on a mixed genetic background, models skin abnormalities, neurological defects, facial dysmorphism, psychomotor retardation, and fertility problems with variable expressivity and incomplete penetrance. Our findings suggest that hemizygosity for *SNAP29* contributes to subset of pathologies associated with 22q11.2DS, including: dysmorphic facies, rib abnormalities, global developmental delays, motor deficits (McDonald-McGinn et al., 2013, Bassett et al., 2011, Oskarsdottir et al., 2005, Roizen et al., 2010, Van Aken et al., 2007), and reveals a role for *SNAP29* in male fertility.

## Methods

### Animals

All procedures and experiments were performed according to the guidelines of the Canadian Council on Animal Care and approved by the Animal Care Committee of the RI-MUHC. CD1 mice were purchased from Charles River laboratories. *Snap29* mutant mouse lines (*Snap29*^*lam/lam*^) were generated on a mixed genetic background (CD1; FvB) and maintained on the outbred CD1 genetic background.

### Generation of Snap29 in situ Probe

An *in situ* probe for *Snap29* was generated from cDNA of wildtype E10.5 CD1 embryos. Briefly, RNA was extracted from embryos using Trizol (Invitrogen) according to the manufacturer’s instructions. SuperScript® II Reverse Transcriptase (Thermo Fisher Scientific) Kit was used to synthesize a complementary DNA (cDNA). Primers were designed and used to amplify exons 2-5, including 207 bp of the 3’ UTR of *Snap29.* Primers used: forward sequence: AGCCCAACAGCAGATTGAAA and reverse sequence: AAAACTCAGCAGAACAGCTCAA. The cDNA fragment was cloned into TOPO using a TA Cloning Kit (Invitrogen). The cloned *Snap29* cDNA was verified by Sanger sequencing. DIG RNA Labeling Mix (Roche) was used to produce digoxigenin labeled sense and antisense probes, according to the manufacturer’s instructions. Briefly, the plasmid was linearized using either XbaI and Kpn1 restriction enzymes (NEB), and SP6 and T7 polymerases (NEB) were used to produce sense and antisense probes, respectively.

### *In situ* hybridization

E7.5-E12.5 wild type embryos were collected from pregnant CD1 females, fixed in 4% paraformaldehyde overnight at 4°C and dehydrated in methanol (for whole mount) or ethanol (for *in situ* sections). For *in situ* hybridization on sections, deciduas and embryos were serially sectioned at 5 µm. Antisense probe was used to detect the expression of *Snap29*, and sense probe was used as a control. All protocols used for whole mount or section *in situ* hybridization were previously described (Revil and Jerome-Majewska, 2013).

### Generation of Snap29 knockout mice line using CRISPR/Cas9

Two *Snap29* mutant mouse lines (*Snap29*^*lam1*^ *and Snap29*^*lam2*^) were generated on a mixed genetic background (CD1 and FvB) using CRISPR/Cas9 methodology (Henao-Mejia et al., 2016). Briefly, four single guide RNA (sgRNA) sequences flanking exon 2 of the mouse *Snap29* gene, (sgRNA1 and sgRNA2 located in intron 1, and sgRNA3 and sgRNA4 located in intron 2) were designed using the online services of Massachusetts Institute of Technology (http://crispr.mit.edu). SgRNAs were synthesized using GeneArt Precision gRNA Synthesis Kit (Thermo Fisher Scientific). 50ng/ul Cas9 mRNA (Sigma) together with 4 sgRNAs (6.25ng/µl) were microinjected into the pronucleus of fertilized eggs collected from mating of wild type CD1 and FvB mice. Injected embryos were transferred into uterus of pseudo-pregnant foster mothers (CD1, Charles River laboratories). All mutant mouse lines were maintained on the mixed CD1 genetic background.

### Genotyping

Yolk sacs were used for genotyping embryos and tail tips were used for genotyping pups from the *Snap29* colony. Standard polymerase chain reaction (PCR) was used for genotyping. Three-primers PCR (primers sequences: Right primer: GACTGAGTCTCACCTGGTCC, Left primer: TGGCTTTTGGAATGACTTG, was optimized to enable detection of embryos and pups carrying wild type (435 bp amplicon) or mutant alleles (240 and 300 bp amplicons, *Snap29*^*lam1*^ *and Snap29*^*lam2*^, respectively) (Supplemental Figure 3). All PCR products were confirmed by Sanger Sequencing.

### Embryo, Brain and Testis collection

Wild type CD1 embryos were used for all *in situ* hybridization experiments. For all other analysis, embryos were collected from *Snap29* mutants on a mixed genetic background (CD1; FvB). For embryo collection, the day of plug was used to indicate pregnancy and designated as embryonic day (E) 0.5. Homozygous mutant embryos and pups were generated from mating of *Snap29* heterozygous mice. For perinatal and adult analysis, the day of birth was designated as postnatal day (P) 0. Embryos and tissues (brain and skin) were collected in RIPA for western blot analysis or fixed in 4% paraformaldehyde and processed as previously described (Revil and Jerome-Majewska, 2013). For testis collection wild type or *Snap29* homozygous mutant male mice were euthanized and testes were dissected, weighed, and fixed immediately with Bouin fixative for 24 hours (hr) before processing and embedding in paraffin blocks as previously described (Russell et al., 1990)

### Western blot analysis

Western blot analysis was performed as previously described (Gupta et al., 2016, Jerome-Majewska et al., 2010). Briefly, P1 dorsal skin were collected in 1× RIPA lysis buffer (25mM Tris-HCl pH7.6, 10% glycerol, 420 mM NaCl, 2mM MgCl2, 0.5% NP-40, 0.5% Triton X-100, 1mM EDTA, protease inhibitor) on ice. Skin samples were then sonicated and centrifuged at 13000rpm for 20 minutes at 4°C. Clarified protein lysates were measured according to standard methods using a DC protein assay kit (Bio-rad, Mississauga, Ontario, Canada). Bradford assays (Bio-Rad) were used to standardize amount of total protein loaded on a gel 12% polyacrylamide gel was then transferred to polyvenyidene fluoride (PVDF membrane, Bio-Rad). 5% nonfat dry milk in PBST (Phosphate buffered saline with Tween 20) was used to block membranes followed by incubation with primary and secondary antibodies. The ECL Western Blotting Detection System (ZmTech scientifique) was utilized to detect the immunoreactive bands and images were taken with Bio-Rad’s ChemiDoc MP System (catalog# 1708280). Primary antibodies used for western blots were rabbit anti-SNAP29 monoclonal antibody (1:5000, Abcam) and rabbit anti-Beta-actin monoclonal antibody (1:5000, Cell Signaling). An anti-rabbit secondary antibody (1:5000, Cell Signaling) was used for both SNAP29 and beta-actin. The bands were quantified by Image lab (Bio-Rad). Statistical significance was tested using Prism 5 (GraphPadVersion 1.0.0.0).

### Immunohistochemistry and Histological Staining of Brain and Eye

Paraffin sections (E17.5, coronal, 6 µm) were used for immunohistochemistry as previously described with slight modification (Lim et al., 2010). Briefly, after deparaffinization and rehydration, the sections were incubated with 1% H202/methanol for 10 min. Antigen retrieval was performed by heating sections at 100°C in 0.01M citric acid for 10min. Sections were blocked with 10% goat serum for 1 hr, incubated with primary antibodies at 4°C overnight, then incubated with appropriate secondary antibody conjugated with biotin (1:1000, Vector Lab) at RT for 1 hr, and incubated with ABC kit (1:1000, Vector Laboratories, PK-6100) at RT for 1 hr. For nuclear staining, sections were treated with 0.5% Triton X-100 for 10 min before the blocking step. The signal was detected with immPACT DAB (Vector Laboratories, SK-4105) and each section was counterstained with diluted Hematoxylin (1:30, Leica Biosystems, 3801570). The primary antibodies used in this study are TBR1 (rabbit, 1:1000, Abcam, ab31940), CTIP2 (rat, 1:200, Abcam, ab18465), Reelin (mouse, 1:200, Millipore, MAB5364), and SATB2 (mouse, 1:200, Bio Matrix Research, BMR00263).

Coronal sections (5 µm) of E17.5 wild type and homozygous *Snap29* mouse embryo brains were stained with cresyl violet in histopathology core of MUHC (McGill University Health Center).

For eye histological section, mice were sacrificed following the ERG recording, and the eyes were removed and fixed for 2 hrs in 4% paraformaldehyde. After a dissection to remove the cornea and lens, the eye cups were placed in 4% paraformaldehyde and fixed overnight on a shaker. On the following day, the eyecups were dyed with 1% osmium tetroxide for 3 hours and subsequently dehydrated with 50%, 80%, 90%, 95% and 100% ethanol. Finally, the eye cups were embedded in resin (Durcupan ACM Fluka epoxy resin kit, Sigma-Aldrich, Canada) and placed in an oven at 55 °C for 48 hrs. The resin embedded eye blocks were then cut into 1.0 μm thick sections with an ultramicrotome (Leica EM UC6 microtome, Leica microsystem, USA) and dyed with 0.1% toluidine blue. Retinal photographs were taken using a Zeiss Axiophot (Zeiss microscope, Germany) at a 40X magnification. The thickness of each retinal layer was measured in the supero-temporal region at a position between 680 μm and 1020 μm from the optic nerve head (Polosa et al., 2017).

### Transmission Electron Microscopy (TEM)

P1 pups were anesthetized and perfused with 2.5% glutaraldehyde in 0.1M sodium cacodylate buffer. 1-mm^3^ dorsal skin section was taken from the perfused pups and placed in the same fixative for an additional 2 hrs at 4°C. The skin samples were rinsed three times in 0.2 M sucrose/ sodium cacodylate buffer and was left in this buffer overnight. The samples were then post-fixed in ferrocyanide-reduced osmium tetroxide for 1 hr at 4°C followed by dehydration with acetone, infiltration with acetone/epon, and embedded in Epon. Sections were cut with Leica Microsystems Ultracut UCT. To examine the sections, FEI Tecnai 12 BioTwin 120 kV Transmission Electron Microscope imaged with an AMT XR80C CCD Camera System.

### Cell counting

Two serial sections from each E17.5 wildtype (n=3) or *Snap29* homozygous mutant (n=4) embryo were used for quantification of each cortical layer marker (total 6 sections for wildtype and 8 sections for mutant). 10x images were taken from the cortical sections (at the level of internal capsule), and a selected area (200 µm wide) from each image as shown in figure 8 with a grid used for cell counting. Each selected area is divided into 8 sub areas (divided with uniformly spaced lines horizontal to the ventricular surface) spanning from the pial surface to the intermediate zone (Grids 1-8; 1 being closest to the pial surface) excluding the ventricular zone and subventricular zone. TBR1, CTIP2, SATB2 positive cells were counted using Automatic Threshold (Yen) in Image J and Reelin positive cells were counted using Manual Threshold (Yen, set value: 215). Total cells (Hematoxylin positive) were counted using Automatic Local Threshold (Otsu, Radius:10). The percentage of TBR1+/total, CTIP2+/total, or SATB2+/total cells was plotted for each 8 Grid for wildtype or mutant. Error bars are Mean +/- SEM. The percentage of Reelin/total cells was plotted for Grid 1 (Error bars are Mean +/- SD). Multiple pairwise t tests were performed in Prism 8. The differences between wildtype and mutants are statistically not significant for all the layer markers tested here.

### BrdU and EdU labeling and staining

Pregnant mice were injected with BrdU (50 mg/kg, i.p.) at E12.5 and E13.5, and with Edu injection (50 mg/kg, i.p.) at E14.5 and E15.5, respectively. Embryos were collected at E17.5 and fixed in 4%PFA for 48 hours. Paraffin sections (E17.5, coronal, 6 µm) were used for BrdU and EdU double staining. After deparaffinization and rehydration, antigen retrieval was performed as described above. The sections were first treated with 2N HCl for 30min at RT, then with 0.1M sodium tetraborate for 10 min, and finally with 0.5% Triton X-100 for 10 min. After blocking with 10% goat serum for 1 hr, primary antibody, anti-BrdU antibody (Abcam, ab6326, 1:100 in 1% goat serum/PBS), was incubated for 4°C overnight on each section. Goat anti-rat IgG-Alexa Fluor 594 (1:200, Invitrogen) was applied as a secondary antibody for 1hr, followed by Click-iT reaction cocktail (Alexa Fluor 488, Invitrogen, C10337) for 30 min according to manufacturer’s recommendation. Hoechst 33342 was used for nuclear staining.

### Electroretinogram (ERG) recordings

After an overnight dark adaptation, flash ERGs were recorded as previously reported (Polosa et al., 2017). Briefly, the mice were anesthetized with an intramuscular injection of ketamine (75 mg /kg) and xylazine (12.5 mg/kg) solution. One drop of 1% Mydiacyl and 0.5% Alcaine were used respectively to dilate the pupil and to decrease blinking. The mice were then placed on their right side on a homeothermic heating pad (Harvard Apparatus, Holliston, MA) at a fixed temperature of 37 ° C in a recording chamber of our design. ERGs were recorded by placing a DTL electrode fiber (27/7 X-Static® silver-coated conductive nylon thread, Sauquoit Industries, Scranton, PA, USA) on the cornea and maintained in position with a coat of moisturizing gel (Tear-Gel Novartis Ophthalmic, Novartis Pharmaceuticals Inc., Canada). A reference electrode (Grass E5 disc electrode) was placed under the tongue of the mouse and a ground electrode (Grass E2 subdermal electrode) was inserted subcutaneously in the tail. Full-field ERGs (bandwidth: 1-1000Hz, 10,000X, 6db attenuation, Grass P-511 amplifiers) were recorded using the Biopac Data Acquisition System (Biopac MP 100WS, Biopac System Inc., Goleta, CA, USA). Scotopic ERGs were evoked to gradually brighter flashes of white light ranging from −6.3 log cds.m^-2^ to 0.9 log cds.m^-2^ in 0.3 log-unit intervals (Grass PS-22 photostimulator, Grass Technologies, Warwick, RI, USA; average of 5 flashes, interval of stimuli: 10sec). Photopic ERGs (background light: 30 cd.m^-2^ flash intensity: 0.9 cds.m^-2^; average of 20 flashes at 1 flash per second) were recorded after a light adaptation period of 20 minutes following the dark-adaptation period. The amplitude of the a-wave was measured from the level of the baseline to the most negative trough while the amplitude of the b-wave is measured from the trough of a-wave to the fourth positive peak.

### Skeletal preparations

Skeletal preparations with Alcian blue to stain cartilage or Alcian blue/Alizarin red to stain cartilages and skeletons of E14.5, E16.5, P1 and P3 embryo and pups were performed as previously described (Hogan B., 1994). Briefly, whole mount E14.5, E16.5 embryos and P1 and P3 pups were treated with ethanol and acetone for 24 hours after eviscerating and removal of skin (for E16.5, P1 and P3 samples) and stained with Alcian Blue/Alizarin Red stain (Sigma) for 3-4 days at 37°C on a rocker. Stained embryos and pups were incubated in 1% KOH for 72-96 hrs and washed with 1% KOH/glycerol mixture.

### Rotarod

Rotarod (47600, KYS Technology, UGO Basile S.R.L, Italy) testing was performed as previously described (Dorninger et al., 2017). Briefly, mice were tested on an accelerating rotarod mode (4-40 rpm) for 300 sec. The latency to fall (in seconds) was recorded. Two rotations in the cylinder were considered as a fall. Each mouse had 3 days training with 3 trials per day and the fourth day was the experimental day. The mean value of the three trials on the experimental day was used for statistical analysis.

### CatWalk Automated Quantitative Gait Analysis

CatWalk program (CatWalk XT 10.6, Noldus, Leesburg, VA. USA) was used to analyze the gait of the mice according to manufactures instructions and published procedures (Hamers et al., 2001). Animals were trained for 3 days before the final measurements were collected. A minimum of 3 compliant runs were acquired with following run Criteria: Min Duration: 0.5 sec, Max Duration: 8 secs, Max Variation: 60%, Min Number of Compliant Runs: 3. Parameters of acquisition are as following: Camera Gain: 15 db, Green Intensity Threshold: 0.3, Red Ceiling Light: 17.4 V, Green Walkway Light: 16 V, Camera Position: 24 cm from the glass. Mean of all compliant runs per mouse were used to calculate gate of mice within different groups, according to sex and genotype. Data acquired on the experimental day were compared for each genotype. All five categories of gait parameters classified in (Hamers et al., 2006) (parameters related to individual paws, the position of footprints, and time dependent relationship between footprints) were assessed. Parameters showing differences on the experimental day were also compared for each training day (day1-3). To include data for different trial days, the average of total measurements (wildtype, heterozygous and mutant) for each parameter on a given day was acquired. Each measurement was then divided by the experimental average and the normalized data from different experiments were pooled together before statistical analysis. All final results indicate parameters that showed a significant difference over all 4 days.

### Grips strength measurement

Mice were held by the base of their tail and lowered toward the mesh of the grip strength meter (Bioseb Model GS3). After grasping with their forepaws, the body of the mouse was lowered to be at a 45-degree position with the mesh. The mouse was then pulled by the tail away from the mesh until the grip was broken. Similarly, after grasping with the hind paws, the mice were lowered at the 45-degree position with the bar and pulled until the grip was broken. Nine trials were performed for each mouse and the average was used as the grip strength score for that mouse (Tanaka et al., 2018).

### Statistical analysis

To assess statistical significance of the different parameters used in this study, we first determined if the data for a given parameter was normally distributed using the D’Agostino and Pearson normality test. We performed ANOVA analysis on normally distributed data followed by the Tukey post-hoc analysis to determine which pair of data was statistically different. If the data did not pass the normality test, we performed a Kruskal-Wallis test followed by a Dunn’s post hoc test to determine which pair of data was statistically different. The p-values of the statistically significant data are represented by *; *≤0.01, **≤0.001 and ***≤0.0001.

## Results

### Snap29 is ubiquitously expressed during mouse embryogenesis

Given that mutations in *SNAP29* are associated with skin and neurological abnormalities in CEDNIK syndrome (Fuchs-Telem et al., 2011, Hsu et al., 2017, Sprecher et al., 2005), we predicted that it would be specifically expressed in ectoderm and ectodermal derivatives. To examine the expression pattern of *Snap29* at earlier stages of embryogenesis, from E9.5 – E12.5, digoxigenin labeled RNA probes were generated and used for *in situ* hybridization. From E9.5 onwards, ubiquitous expression of *Snap29* was found by wholemount (Supplemental Figure 1) and section (Figure 1A) *in situ* hybridization. Thus, *Snap29* was expressed in derivatives of all three germ-layer, although patients with mutation in this gene show mostly neurocutaneous abnormalities.

**Figure 1.**
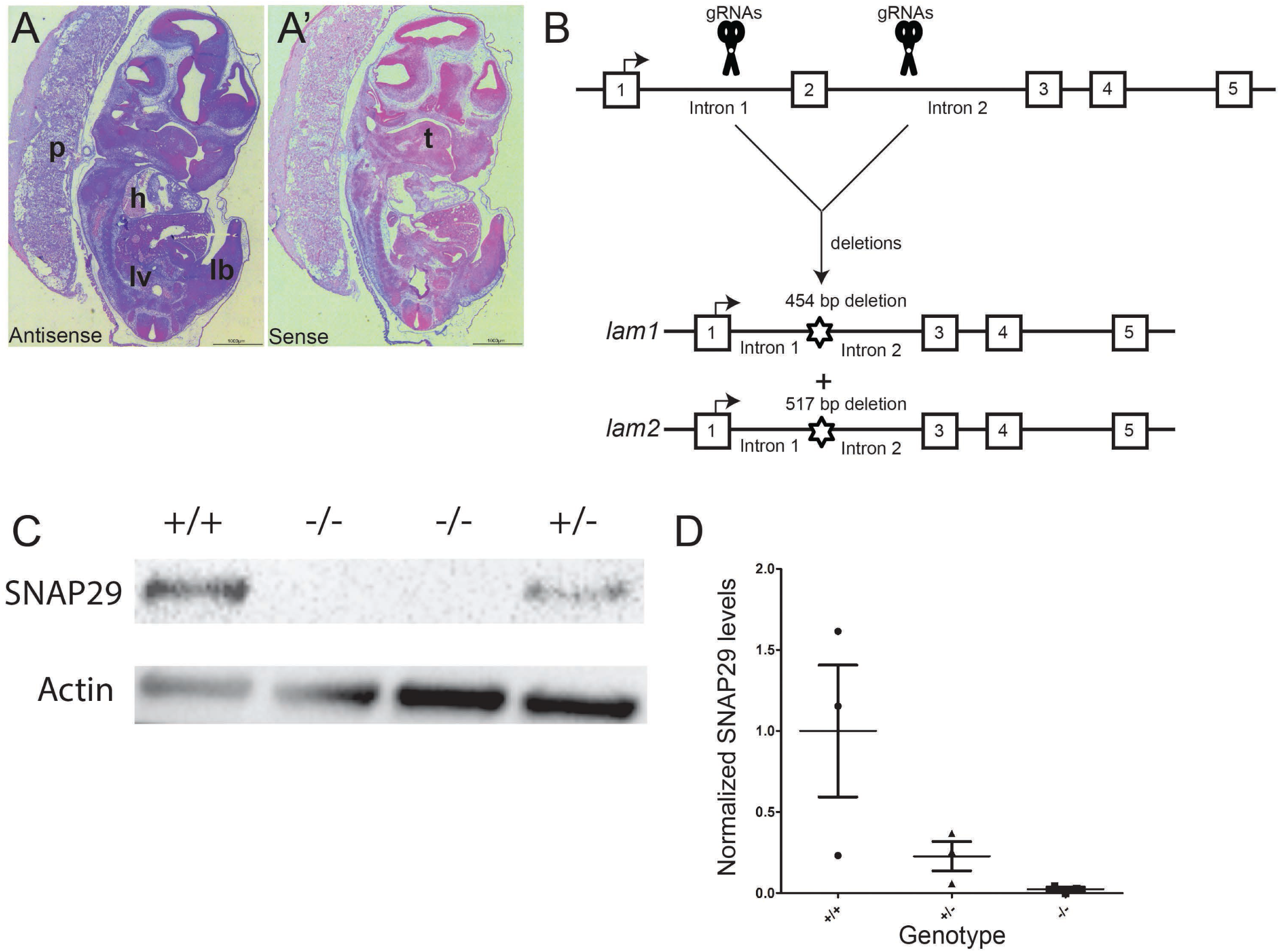
*Snap29* mRNA is ubiquitously expressed and CRISPR-mediated targeting of exon 2 depletes SNAP29 protein. A-A ‘. In situ hybridization with an antisense mRNA probe to *Snap29* reveals ubiquitous expression mRNA expression at E12.5 (Blue) and faint to no signal with a sense control. B. CRISPR design and the resulting targeted alleles, one with a 454bp and one with a 517 bp deletion. C. Western blot analysis of dorsal skin preps of P1 pups either wild-type, heterozygous or homozygous mutant for the *Snap29* allele. D. Quantification of SNAP 29 expression. SNAP29 levels are normalized to WT. Lb: limb bud, l: lung, hrt: heart, s: somite, tb: tail bud. In Panel A and A’ Nuclei are stained pink with Nuclear Fast Red.

### Generation of Snap29 mutant mouse line on a mixed genetic background

Constitutive loss of function mutation of *Snap29* on the inbred C57Bl/6 genetic background results in neonatal death due to severe skin abnormalities and barrier defects (Williams et al., 2016, Schiller et al., 2016a). Since non-epidermal abnormalities associated with CEDNIK syndrome were not described on the C57 inbred genetic background, we reasoned that a mouse model with loss of function mutation of *Snap29* on a mixed/outbred genetic background would model most abnormalities found in human patients. To delete exon 2 of *Snap29* and generate a frameshift and premature termination signal in this gene (Schiller et al., 2016a), we microinjected CRISPR/Cas9 and gRNAs flanking exon 2 of *Snap29* into mouse zygotes on a mixed genetic background (CD1; FvB) (Figure 1B). Of 14 mice born, four carried Sanger-sequence confirmed deletions of exon 2 as well as 267bp and 320bp of flanking intronic sequences resulting in two mutant mouse lines (*Snap29*^*lam1*^ and *Snap29*^*lam2*^). Females carrying deletions of exon 2 were mated to wild type CD1 males to establish *Snap29* mutant mouse colonies. However, since our analysis of *Snap29*^*lam1*^ and *Snap29*^*lam2*^ mutant embryos and pups revealed similar penetrance and expressivity in the phenotypes described below, after 6-generation of back-crossing to the outbred CD1 genetic background, these numbers have been combined.

### Mice with loss of function mutation of Snap29 on a mixed genetic background (CD1; FvB) survive to adulthood

*Snap29* heterozygous mutant mice showed no apparent morphological abnormalities, survived to adulthood and were fertile. *Inter se* mating of *Snap29* heterozygous mice revealed normal Mendelian segregation of *Snap29* mutant alleles at mid-gestation and at birth (Table 1). Furthermore, although 35% of *Snap29* homozygous mutant pups died within the first few days of life (n=14 of 40), the majority, 65%, survived to weaning and adulthood (n= 26 of 40; Table 2). Using western blot analysis, we showed that SNAP9 protein was reduced in dorsal skin of heterozygous P1 pups and undetectable in homozygous mutant samples (Figure 1C and D, Supplemental figure 2 and 3) when compared to wildtype controls, confirming that deletion of exon 2 abolishes SNAP29 protein production. Thus, we concluded that genetic modifiers on the mixed genetic background allow mice with loss of function mutation in *Snap29* to survive to adulthood.

**Table 1:** Number of embryos or pups per *Snap29* genotype

| stage | +/+ (dead) | +/- (dead) | -/- (dead) | # litters |
| --- | --- | --- | --- | --- |
| E11.5 | 28 (0) | 45 (0) | 29 (0) | 9 |
| E16.5 | 36 (0) | 51 (0) | 33 (0) | 10 |
| P1 | 37 (0) | 52 (0) | 39 (0) | 12 |

**Table 2:** *Snap29* skin abnormalities

| Genotype | none (died) | mild (died) | moderate (died) | severe (died) |
| --- | --- | --- | --- | --- |
| -/- | 7 (2) | 14 (1) | 14 (8) | 5 (3) |
| +/- | 0 | 0 | 0 | 0 |
| +/+ | 0 | 0 | 0 | 0 |

**Figure 2.**
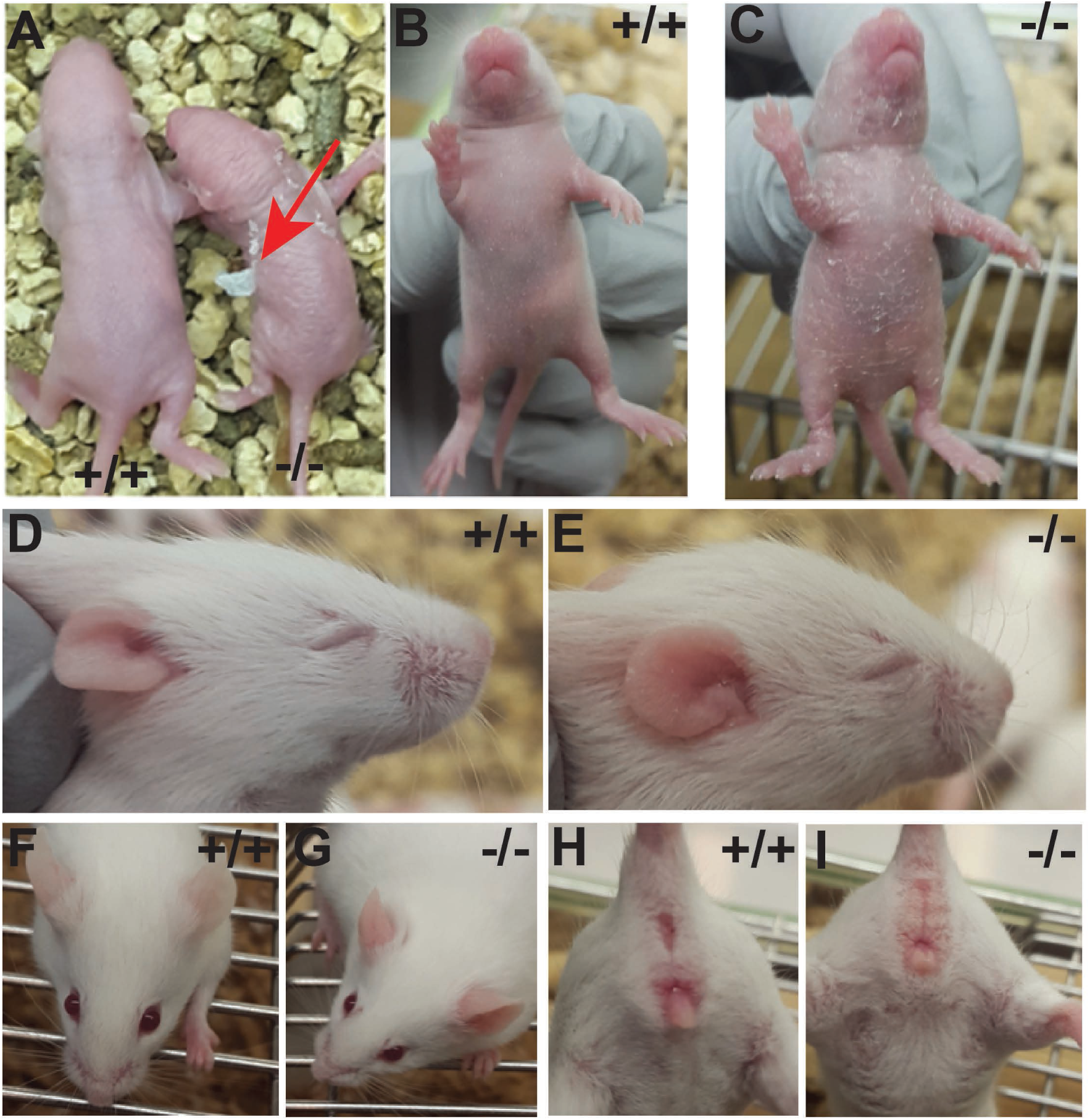
Snap29 mutant mice show diverse skin defects. A. Example of the severe skin peeling observed in *Snap29* ^-/-^ P2 pup (red arrow). B-C. P3 pups showing mild skin peeling. D-E. P11 pups showing an example of nose icthyosis. F-G. Example of reddish and thickened ears in P30 pups. H-I. Example of swollen reddish genetalia.

### Epidermal defects in Snap29 homozygous mutant pups

We next assessed whether *Snap29* homozygous mutant pups and adult mice exhibited pathologies and abnormalities found in CEDNIK and 22q11.2DS patients. We followed 40 *Snap29* homozygous mutant pups born from mating between *Snap29* heterozygous mice, from birth until weaning (n=18 litters). A small fraction of these homozygous mutant pups died within a few days of birth with no apparent defects (n=2 of 40). However, 86% of homozygous mutant pups (n= 33 of 38) developed skin defects between P2 and P6. These abnormalities were classified as severe, moderate or mild depending on the extent of scaling or peeling observed (Figure 2 and Table 2). Most, of the skin defects found in homozygous mutants were in the moderate to severe category (n=19 of 33). In addition, though a subset of homozygous mutant pups with skin abnormalities died within the first few 7-days of life (n=12 of 33), the majority survived (n=21). Furthermore, though Schiller et al reported reduced hair follicles in keratinocyte-specific *SNAP29* mutant embryos on the C57 genetic background (Schiller et al., 2016b) we found that surviving homozygous mutants on the mixed genetic background recovered from their skin defects and form coats that were undistinguishable from their litter mates. Nonetheless by weaning, 100% of surviving homozygous mutants had thickened and reddish ears, scaling on their ears and paws (Figure 2E, 2G and data not shown), and swollen reddish genitalia (Figure 2I), including those with no obvious skin defects in the early perinatal period (n= 5). Moreover, a few homozygous mutants showed severe ichthyosis on their nose later in life (n =3/33, data not shown). Thus, *Snap29* homozygous mutant mice on a mix genetic background recapitulate variable expressivity of skin abnormalities found in CEDNIK patient.

To determine the basis of skin defects in *Snap29* homozygous mutant pups, histological analysis of dorsal skin was performed. Hematoxylin and Eosin (H&E) staining at E16.5 revealed no morphological differences between wild type, *Snap29* heterozygous and homozygous mutant samples (data not shown). However, at P1-prior to onset of skin abnormalities and P3 – when skin abnormalities were morphologically apparent, epidermis of *Snap29* homozygous mutant pups showed hyperkeratosis, and condensed stratum corneum (Figure 3A-C). In addition to the disrupted stratum corneum (Figure 4D and E), transmission electron microscopy of P1 dorsal skin also revealed remnants of organelles in the lower layers of the stratum granulosum (Figure 3F-G), similar to those previously reported by Schiller et al (Schiller et al., 2016a). Thus, epidermal structure is disrupted by loss of function mutation of *Snap29* on the mixed genetic background, similar to what was observed on the inbred genetic background.

**Figure 3.**
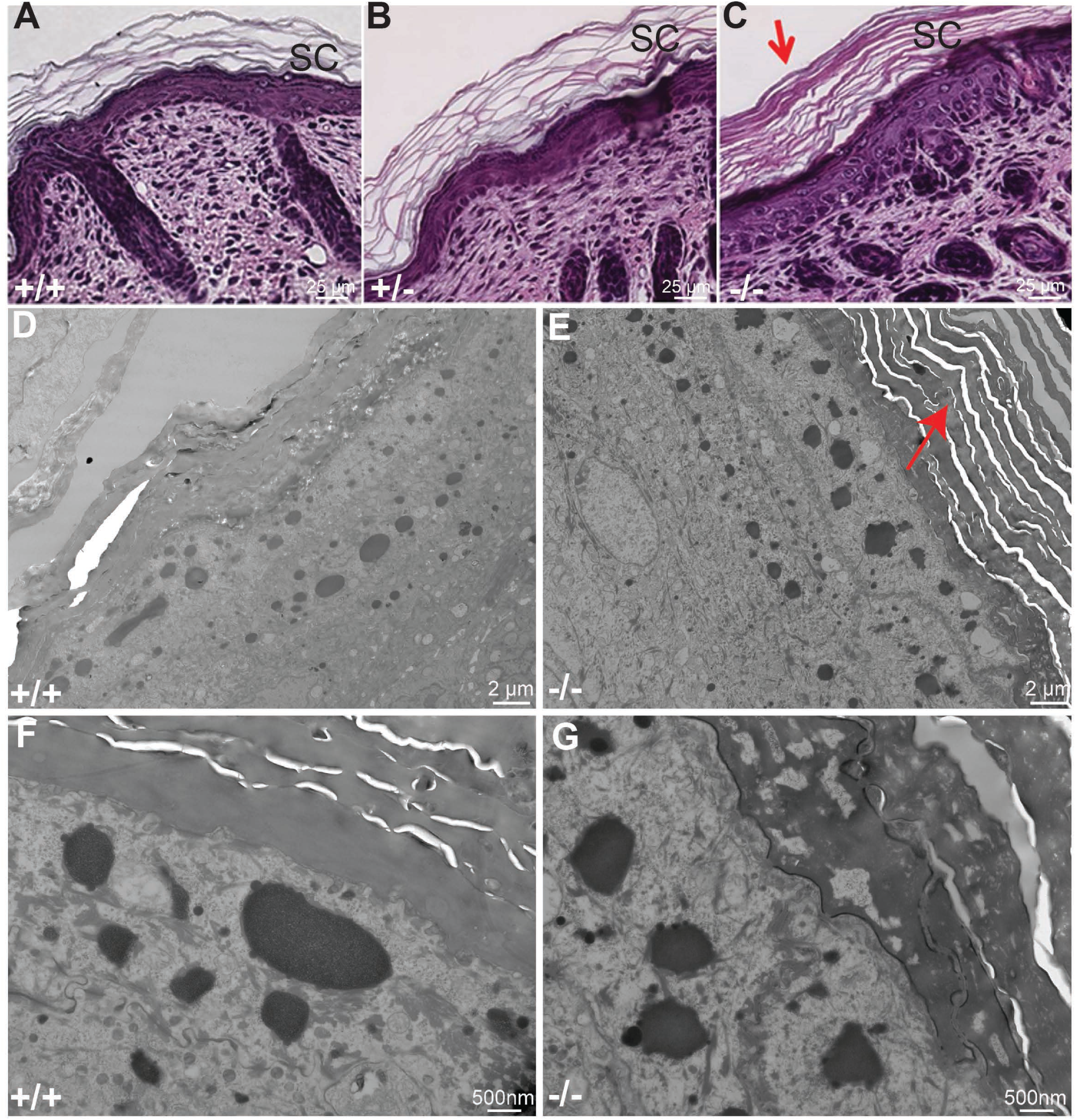
*Snap29* mutant mice show signs of hyperkeratosis. Hematoxylin and eosin staining (A-C) and TEM (D-G) of P1 dorsal skins. The stratum corneum (SC) in *Snap29*^-/-^ mice was found thicker and more condensed (C, E, red arrow) than that of the *Snap29*^*+/+*^ or *Snap29*^*+/-*^ mice.

**Figure 4.**
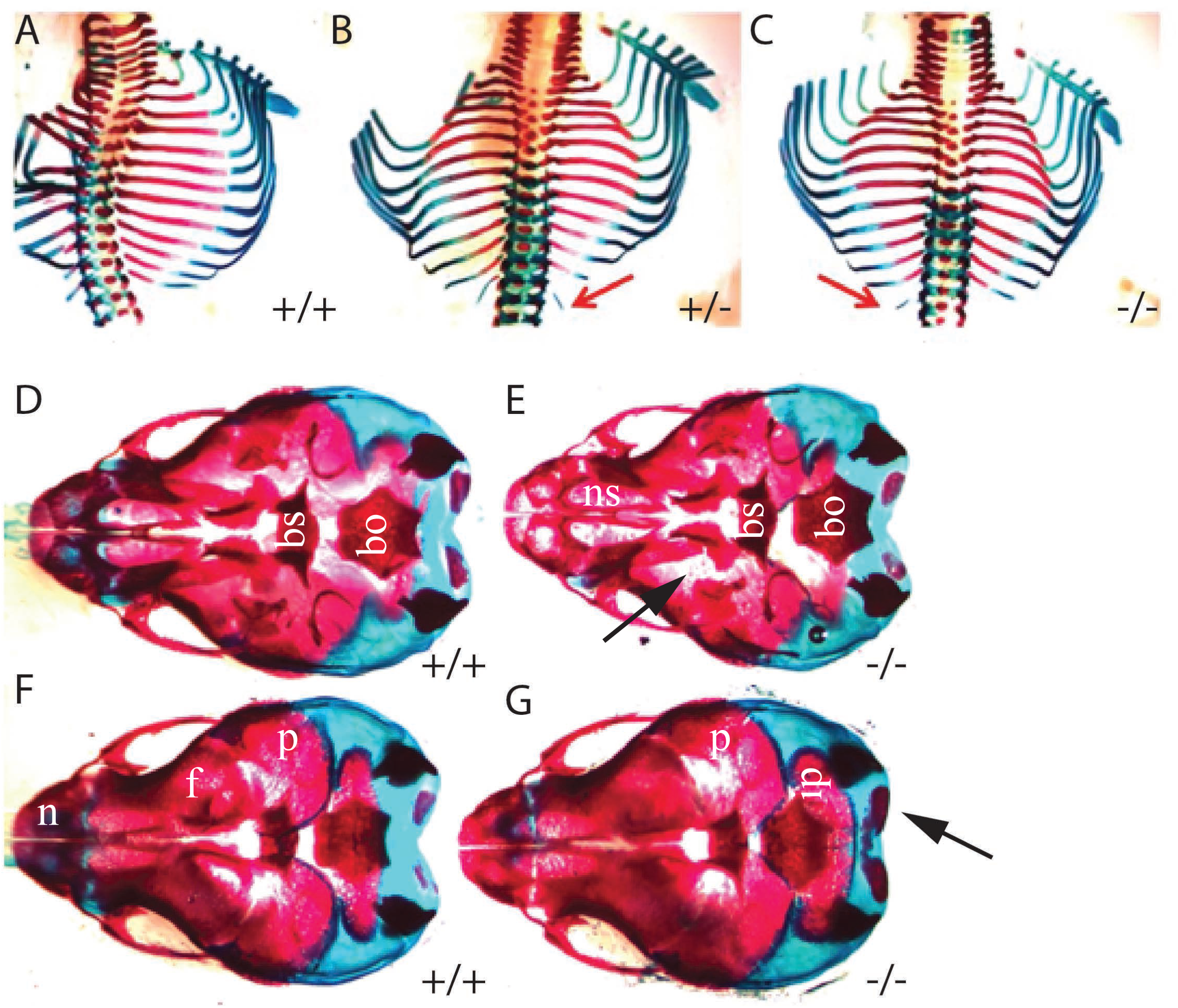
Snap 29 heterozygous and homozygous mutant mice have skeletal defects. Heterozygous (B) and homozygous mutant (C) embryos had extra lumbar ribs (red arrows) that were not seen in control mice (A). Ventral view of heads from *Snap*^*+/+*^ and *Snap29*^-/-^ pups showing reduced mineralization in the head (arrows) (DE). Dorsal view of heads from *Snap*^*+/+*^ and *Snap29*^-/-^ pups indicating increased mineralization of the supraoccipital bone (FG, black arrow) in homozygous mutant embryos. P: parietal, f:frontal, ip: intraperitoneal, bs:baso-sphenoid, ns: nasal capsule, bo; baso-occipital, n:nasal.

Although epidermal defects were found, we surmised that *Snap29* homozygous mutants survived on the mixed genetic background because they were capable of forming a functional skin barrier. To test this hypothesis, an X-Gal skin permeability assay was used to assess the integrity of the epidermal barrier of E17.5 and P1 litters. At E17.5, 30% of wild type (n=3 of 10), 47 % of *Snap29* heterozygous (n= 8 of 18) and 80% of *Snap29* homozygous (n = 12 of 14) mutant embryos, showed X-Gal staining on the ventral surface of the body wall (Supplemental Figure 4), indicating a delay of barrier formation in *Snap29* mutants (Table 3). However, at P1 no X-Gal staining was found in wild type (n=14), heterozygous (n=29) and homozygous mutant pups (n=11) (data not shown). Thus, although skin barrier formation is delayed in *Snap29* homozygous mutant embryos, a proper skin barrier was present by birth. Our data indicates that modifiers on the mixed genetic background rescue the barrier phenotype responsible for neonatal death of *Snap29* homozygous mutant pups on the C57 genetic background.

**Table 3:** Barrier defects in *Snap29* mice

| barrier defects | +/+ | +/- | -/- |
| --- | --- | --- | --- |
| none | 7 | 9 | 3 |
| partial | 3 | 8 | 12 |
| full | 0 | 0 | 0 |

### Skeletal abnormalities in Snap29 pups

CEDNIK and 22q11.2DS patients have skeletal abnormalities as well as dysmorphic facial features (Hsu et al., 2017). To determine if *Snap29* mutants model any of these abnormalities we analyzed cartilage and skeletal development in litters from *inter se* mating *Snap29* heterozygous mice at 4 stages (E14.5, E16.5, P1 and P3). Skeletal abnormalities were found at low penetrance in the face, the head and ribs of *Snap29* mutant embryos and pups. The most common defect found was abnormal mineralization as assessed by staining of bones with alizarin red. Mineralization of the parietal and occipital bones were either delayed (n=3 of 14) or advanced (n=6 of 14) in homozygous mutants (Figure 4 D-G) when compared to wild type (n=11) and *Snap29* heterozygous (n=25) litter mates. In addition, *Snap29* heterozygous (n=8 of 17) and homozygous mutant embryos and mice (n=13 of 32) had extra lumbar ribs (14 pairs of ribs) while only a single such case was seen in one wildtype embryo (n=12) (Figure 4 A-C). Thus, *Snap29* homozygous mutant mice on a mixed genetic background model a subset of skeletal abnormalities and dysmorphisms found in patients with mutations in *SNAP29*, though these phenotypes showed incomplete penetrance and variable expressivity.

### Psychomotor retardation and motor defects in Snap29 homozygous mutant mice

Severe global developmental delay is a hallmark of CEDNIK patients (Fuchs-Telem et al., 2011, Hsu et al., 2017, Sprecher et al., 2005), thus we assessed *Snap29* mutants for motor defects. Newborn *Snap29* homozygous mutant pups appeared to move slower than their wild type and heterozygous litter mates. Therefore, we tested psychomotor function in newborn P1 and P3 (n=3 litter per stage) pups. Briefly, pups were place on a supine position on their back and were timed on how long it took them to move to a prone position – each pup was turned 3 times and average of their turn times were compared between genotypes. This test revealed no significant difference in the ability of pups to move from the supine to the prone position (Supplemental Figure 5). However, we found that a subset of *Snap29* homozygous mutant pups were unable to stand up on their feet after turning onto their stomach (n = 6 of 11) when compared to *Snap29* wildtype (n = 0 of 13) and heterozygous (n= 0 of 21) littermates.

**Figure 5.**
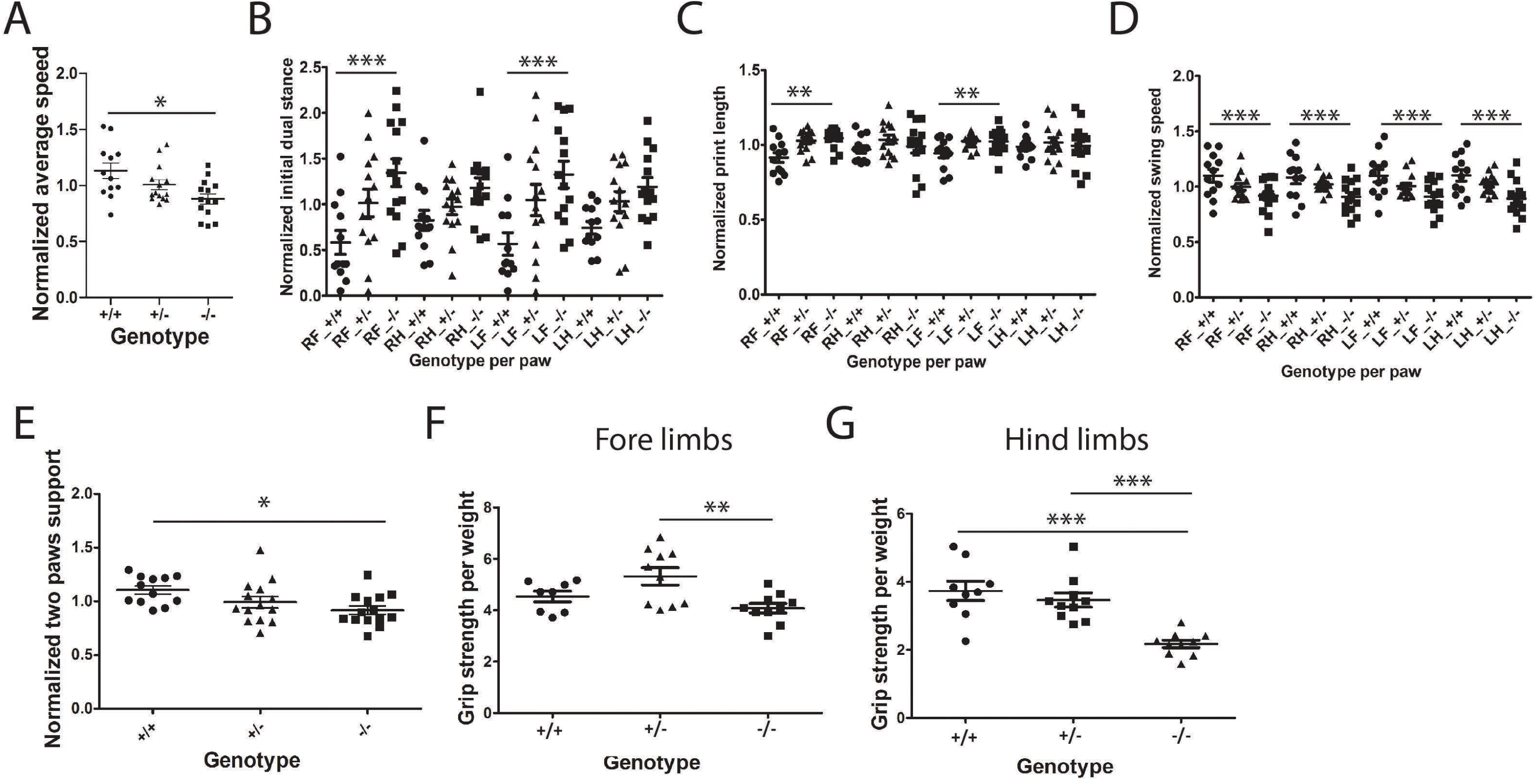
Snap29 mutant female mice exhibit gait defects as measured by Catwalk. The Catwalk system was used to monitor gait parameters in *Snap29*^*+/+*^, *Snap29*^*+/-*^and *Snap29*^-/-^females. The run characteristic parameter run average was significantly slower in *Snap29*^-/-^females (A). The temporal parameter initial dual stance was significantly elevated in both front paws (B). The spatial parameter print length was significantly increased in RF and LF of *Snap29*^-/-^ (C). The kinetic parameter swing speed was decreased in all paws of homozygous mutant animals (D). The interlimb coordination parameter support on two paws was significantly reduced in *Snap29*^-/-^ females (D). Grip strength was assessed for fore limbs (F) and hind limbs (G). *Snap29*^-/-^ females exhibited weaker fore and hind limbs than *Snap29*^*+/+*^ animals. RF: right-front paw; RH: right hind paw; LF: left front paw and LH: left hind paw. Statistical significance: *: p<0.05, **: p<0.01 and ***: p<0.001.

To determine if motor defects persist in adult *Snap29* homozygous mutant mice we used a rotarod, to evaluate neuromuscular coordination, balance and grip strength (Crawley, 1999) in adult males at 5-weeks of age. We found that *Snap29* homozygous mutant male mice (n=3) had a shorter latency to fall, when compared to wild type (n=3) and heterozygous mice (n=3) (Supplemental Figure 6). These findings suggest that *Snap29* homozygous mutant males exhibit deficiencies in neuromuscular coordination, balance and/or grip strength.

**Figure 6.**
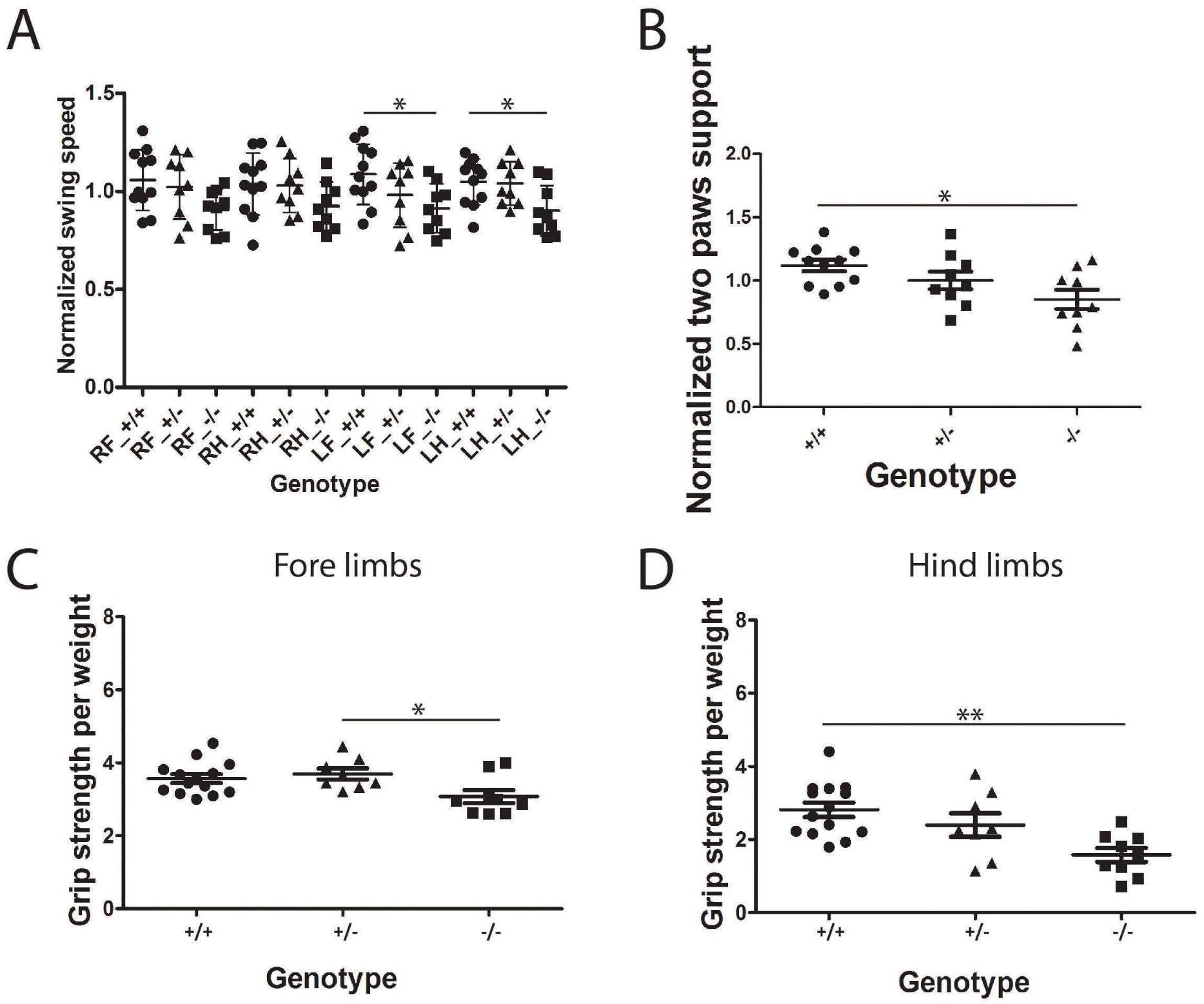
Snap29 mutant male mice exhibit fewer gait defects as measured by Catwalk. Catwalk assay was used to monitor gait parameters in *Snap29*^*+/+*^, *Snap29*^*+/-*^and *Snap29*^-/-^males. The kinetic parameter swing speed was decreased on the left side of *Snap29*^*-/*^*-* males (A). The interlimb coordination parameter support on two paws was significantly reduced in *Snap29*^-/-^ males (B). Grip strength was assessed for fore limbs (C) and hind limbs (D). *Snap29*^-/-^ males exhibited weaker hind limbs than *Snap29*^*+/+*^ animals. RF: right front paw; RH: right hind paw; LF: left front paw and LH: left hind paw. Statistical significance: *: p<0.05, **: p<0.01 and ***: p<0.001.

In parallel, gait, locomotion function and coordination were assessed in 6-weeks old male and female mice using the CatWalk system (Hamers et al., 2006). We found no correlation between weight of mice and average speed run (Supplemental Figure 7), therefore, experimental data were adjusted by normalizing against experimental average for each parameter or for each paw per parameters. Furthermore, as speed can be a confounding factor in these studies, we confirmed that the average speed of all animals used for analysis was at speeds that correlate to a “walk” gait as defined by Bellardita and Kiehn (Bellardita and Kiehn, 2015). The parameters measured by the Catwalk were organized into five major groups: (1) Run characterization, (2) Temporal, (3) Spatial (4) Kinetic and (5) Interlimb coordination, as defined by Gaballero-Garrido et al (Caballero-Garrido et al., 2017) (see Table 4). In all 5 groups significant differences were found when *Snap29* homozygous mutant mice when compared to control littermates.

**Table 4:** Catwalk analysis of Snap29 mutant animals

Catwalk analysis of Snap29 mutant animals
|  | male |  |  |  |  | female |  |  |  |  |  |
| --- | --- | --- | --- | --- | --- | --- | --- | --- | --- | --- | --- |
|  | animal | RF | RH | LF | LH | animal | RF | RH | LF | LH |  |
| average speed | n.s. | n.a. | n.a. | n.a. | n.a. | 0.0185↓ | n.a. | n.a. | n.a. | n.a. | run characterization |
| initial dual stance | n.a. | n.s. | n.s. | n.s. | n.s. | n.a. | 0.0002↑ | n.s. | 0.0002↑ | n.s. | temporal |
| terminal dual stance | n.a. | n.s. | n.s. | n.s. | n.s. | n.a. | <0.0001↑ | n.s. | <0.0001↑ | n.s. |  |
| Print length | n.a. | n.s. | n.s. | n.s. | n.s. | n.a. | 0.0011↑ | n.s. | 0.0011↑ | n.s. | Spatial |
| swing speed | n.a. | n.s. | n.s. | 0.0179↓ | 0.0179↓ | n.a. | 0.0009↓ | 0.0009↓ | 0.0009↓ | 0.0009↓ | Kinetic |
| body speed | n.a. | 0.0129↓ | 0.0129↓ | 0.0129↓ | 0.0129↓ | n.a. | 0.002↓ | 0.002↓ | 0.002↓ | 0.002↓ |  |
| support two paws | 0.019↓ | n.a. | n.a. | n.a. | n.a. | 0.0202↓ | n.a. | n.a. | n.a. | n.a. | Interlimb coordination |
| Support three | 0.0404↑ | n.a. | n.a. | n.a. | n.a. | n.s. | n.a. | n.a. | n.a. | n.a. |  |
| BOS_FrontPaws | n.s. | n.a. | n.a. | n.a. | n.a. | 0.022*↑ | n.a. | n.a. | n.a. | n.a. |  |
| Phasedispersion_LF_LH_Cstat | 0.0172↑ | n.a. | n.a. | n.a. | n.a. | n.s. | n.a. | n.a. | n.a. | n.a. |  |
| Phasedispersion_LF_RH_Cstat | n.s. | n.a. | n.a. | n.a. | n.a. | 0.0297↑ | n.a. | n.a. | n.a. | n.a. |  |
| Couplings_LH_LF_Cstat | 0.0177↓ | n.a. | n.a. | n.a. | n.a. | n.s. | n.a. | n.a. | n.a. | n.a. |  |
| Couplings_LF_LH_Cstat | 0.0256↑ | n.a. | n.a. | n.a. | n.a. | n.s. | n.a. | n.a. | n.a. | n.a. |  |

\* difference is between +/+ vs +/-
n.s.= not significant
n.a.= not applicable
↓=down

**Figure 7.**
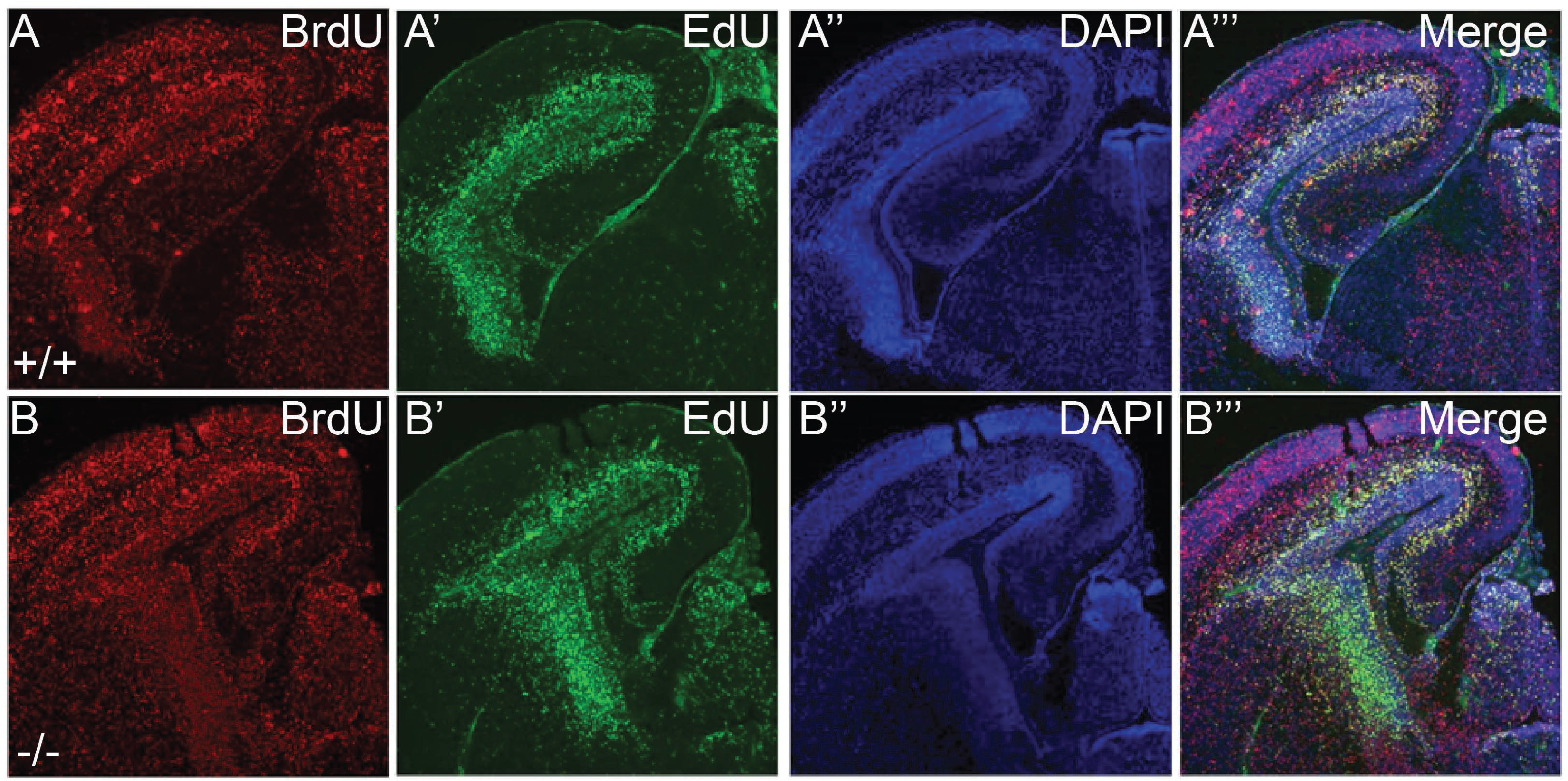
Snap29 E17.5 mutant brains show increased proliferation. Representative images of E17.5 brain of *Snap29*^*+/+*^ (A-A’’’) and *Snap29*^-/-^ (B-B’’’) mice. BrdU is shown in red (A, B), EdU is green (A’-B’), the nuclear marker DAPI in blue (A’’, B’’) and the merge image is shown last (A’’’-B’’’). BrdU was injected first at E13.5 and EdU was injected at E15.5 prior to dissection. We observed an increase in BrdU and in EdU signals in the brain of *Snap29*^-/-^ brain suggesting increased proliferation.

Significant differences were found in all five parameters analyzed from the Catwalk when female *Snap29* homozygous mutants (n=14) were compared to control female littermates (n=12 wildtype and n=14 heterozygous). For run characterization, average speed was significantly reduced in female *Snap29* homozygous mutants, when compared to wild type female litter mates (Figure 5A; p= 0.0185, ANOVA). However, this change was not associated with a significant difference in the number of steps or the duration of runs for these female mice (Table 4). In addition, several parameters in the Step Cycle (time in seconds of ground contact for either fore (F) or hind (H) limb) were significantly affected in *Snap29* homozygous mutant females. Specifically, the Temporal parameters, initial dual stance – time of the initial step in each Step Cycle when both hind or front paws simultaneously make contact with the glass plate (Figure 5 B; p= 0.0002) and terminal dual stance – second step of each Step Cycle (Supplemental Figure 8B; p= <0.0001) were significantly increased in right fore limb (RF) and left fore limb (LF) of female *Snap29* homozygous mutants when compared to female controls. Consistent with the temporal changes, the spatial parameter print length – the distance between two prints was also significantly increased in RF and LF of female homozygous mutants (Figure 5C; p = 0.0011) when compared to controls.

Two kinetic parameters, swing speed, the velocity (distance/time) when paw is not in contact with glass plate (Figure 5D; p=0.0009) and body speed (Supplemental Figure 8A and 8B; p =0.0129) were significantly decreased in all paws of female homozygous mutant mice when compared to controls. In addition, Catwalk parameters associated with interlimb coordination were also significantly different in female *Snap29* homozygous mutants when compared to controls. Specifically, diagonal support on two paws was significantly decreased in female homozygous mutant mice, (Figure 5E; p= 0.0202), and although support on three paws increased in this group, the difference was not statistically significant. In addition, Phase Dispersion LF_right hind limb (RH) Cstat, also referred to as Phase Lag using Circular Statistics, was significantly increased in female homozygous mutants (Supplemental Figure 8D; p=0.0297). Altogether data acquired using the Catwalk indicate that female *Snap29* homozygous mutant mice move slower than controls, and show increased contact with the ground using their forelimbs.

In contrast to what was found in female, differences in run characterization, temporal and spatial parameters in male *Snap29* homozygous mutant males (n=9), were not statistically significant when compared to male controls (n= 11 wild-type, n=9 for heterozygous). However, changes in kinetic parameters and interlimb coordination were significantly different in male homozygous mutants when compared to controls. Thus, velocity was significantly reduced in RF, LF and left hind limb (LH) (Figure 6A; p<0.05) and body speed was significantly decreased in all paws of male homozygous mutant mice when compared to controls (Supplemental figure 9A and 9B; p =0.0129). For interlimb coordination, diagonal support on two paws was significantly decreased (Figure 6B p=0.0202), while support on three paws was significantly increased in male (Supplemental Figure 9B; p= 0.019) when compared to control. Furthermore, in mutant homozygous mutant males, Phase Dispersion LF_LH Cstat, (Supplemental Figure 9C; p=0.0172), and Cstat of couplings LF_LH, (Supplemental Figure 9D; p=0.0177), the temporal relationship between placements of two paws within a Step Cycle, were significantly increased when compared with controls. Consistent with these observations Cstat for the couplings LH_LF was significantly decreased in homozygous mutant males (Supplemental Figure 9E; p=0.0256), when compared to controls. Overall the Catwalk revealed reduced speed in *Snap29* homozygous mutant males and indicate that their fore limbs had increased contact with the ground when compared to controls. These studies suggest that *Snap29* homozygous mutant mice may suffer from weakness or paralysis in their hind limbs.

**Figure 8.**
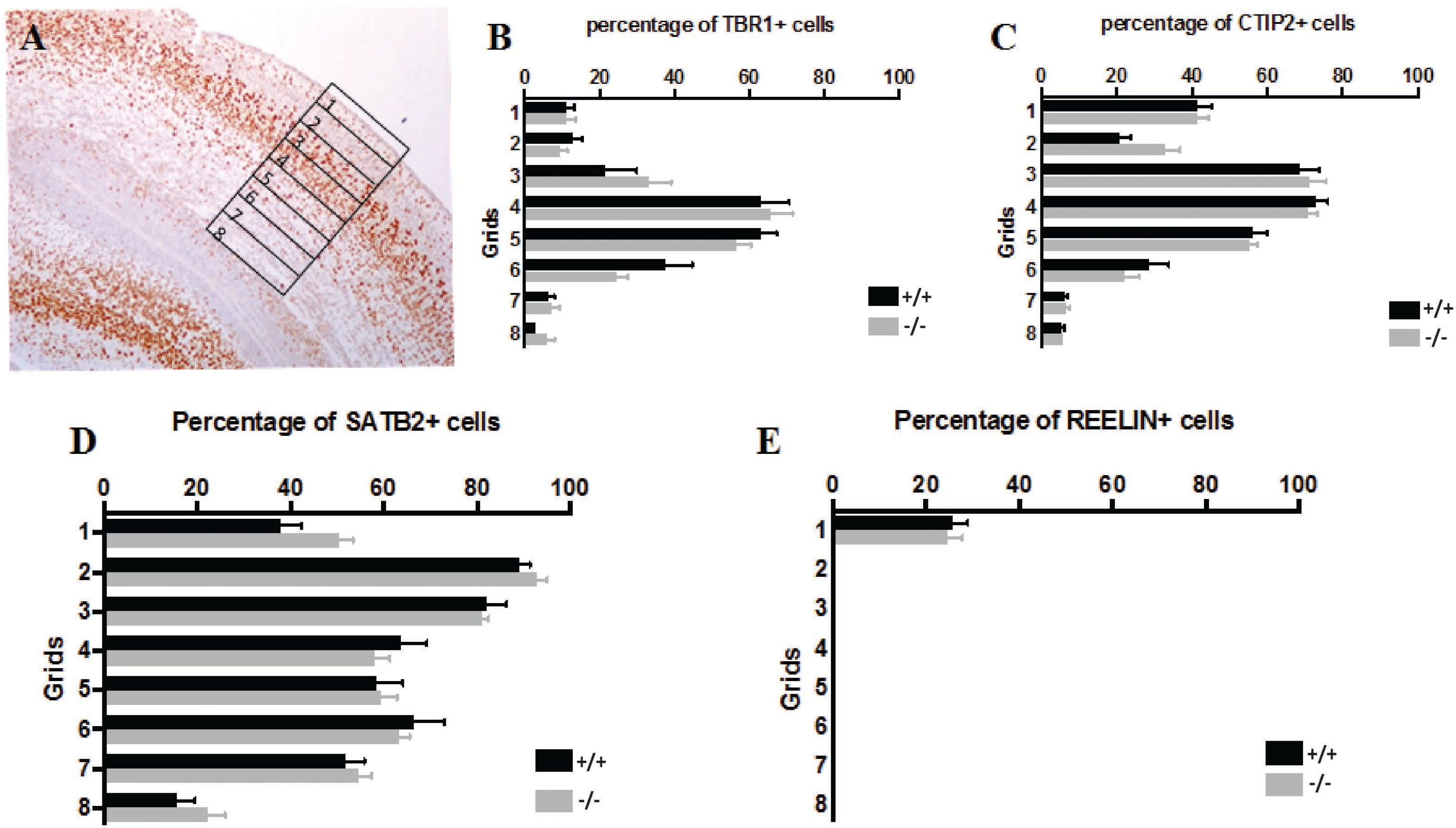
Counting of cortical layer organization markers. A. Representative images showing cell counting. B. Counting of Tbr1; C. Stip2; D. Satb2; E. Reelin stained neurons did not show differences between *Snap29* genotypes. The total number of cells was counted using automatic local threshold.

**Figure 9.**
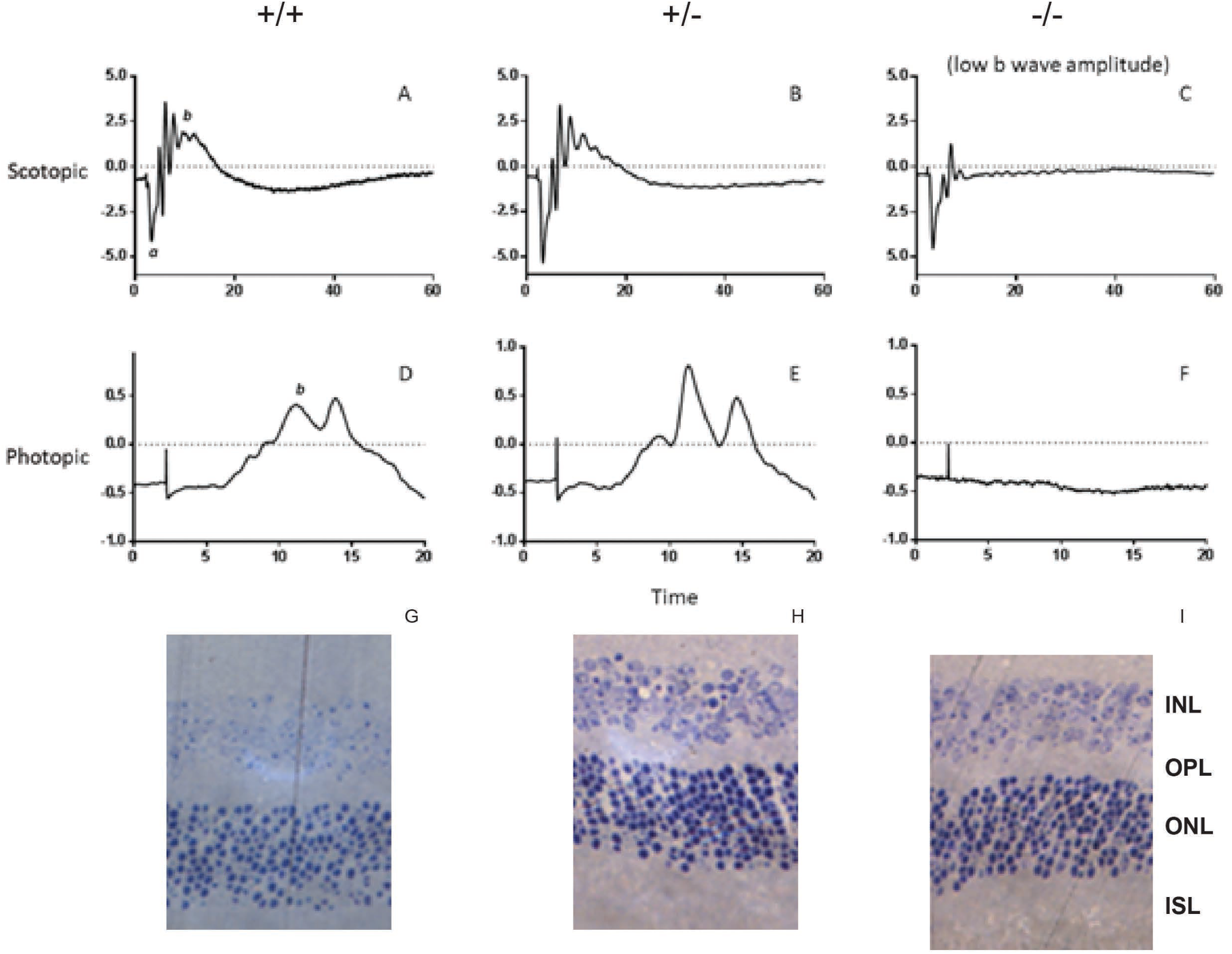
Analysis of ophthalmological defects in *Snap29*^-/-^ mice. A-F. Representative scotopic (A-C) and photopic (D-F) ERG wave forms in wild type (A, D, G), heterozygous (B, E, H) and homozygous mutant mice (C, F, I). A: scotopic ERG recorded from wild type mouse; B: scotopic ERG recorded from heterozygous mouse; C: scotopic ERG recorded from homo-zygous mouse with low b wave amplitude; D: photopic ERG recorded from wild type mouse; E: photopic ERG recorded from heterozygous mouse; F: photopic ERG recorded from homozygous mouse with low b wave amplitude. G-H. Representative eye histology of wild type male mouse (G) at P118; heterozygous female mouse at P83 (H) and homozygous mutant female at P87 (I). ISL: inner segment layer; ONL: outer nuclear layer; OPL: outer plexiform layer; INL: inner nuclear layer.

To test if deficiencies uncovered by the rotarod and Catwalk tests were due to changes in strength, a grip meter was used to measure strength of 7-weeks old male and female mice. Grip strength was significantly reduced in fore limbs of female *Snap29* homozygous mutant mice (Figure 5F; n= 10; p=0.0068) when compared to heterozygous, but not wildtype controls. However, grip strength of hind limbs of female homozygous mutants was significantly decreased when compared to wildtype controls (Figure 5G; p=0.0002). Differences in strength persisted when these animals were retested at 14 weeks of age. But, in addition to reduced grip strength in hind limbs, grip strength was also significantly decreased in fore limbs of homozygous female mice when compared to wild type controls (Supplemental Figure 8E-F; p<0.0124 and 0.0291 respectively). Grip-strength was also reduced in male homozygous mutants. Similar to their female counterparts, significantly reduced fore limb strength was found for *Snap29* homozygous mutant males (p=0.0217; Figure 6C) when compared to heterozygous litter mates (n=10), while hind limb strength was significantly different in homozygous mutant males (Figure 6D; p= 0.0025 when they were compared to wildtype controls (n=9). Altogether these data indicate that *Snap29* homozygous mutant mice present with reduced hindlimb grip strength which progresses to involve the forelimb as the animals aged. Thus hypotonia, similar to the neurological/muscle abnormalities found in CEDNIK patients are found in *Snap29* homozygous mutant male and female mice.

### Normal neurogenesis and lamination of the cerebrum of Snap29 homozygous mutant embryos

Since patients with CEDNIK show severe structural and neurological abnormalities in the brain, and given that *Snap29* mutant mice exhibited motor defects and no skeletal malformations in their limbs (data not shown), we first assessed whether these mice exhibited brain defects. We first tested if neurons were either not produced or were not proliferating in *Snap29* embryonic brain. We assessed proliferation of neurons by injecting pregnant females with BrdU at E13.5 and followed by EdU at E15.5 and confocal microscopy to detect new incorporation of those DNA analogs *in vivo* at E17.5 (Figure 7). We observed increased labelling of BrdU and EdU in one of four *Snap29* homozygous mutant embryos examined (Figure 7B-B’’”), when compared to wild-type siblings (Figure 7A-A’’’). Thus, we surmised that proliferation may be perturbed in a small subset of *Snap29* homozygous mutant mice but with low penetrance.

To determine if changes in proliferation result in perturbed neurogenesis, cortical lamination was examined. Cortical lamination was evaluated using molecular markers in WT and *Snap29* homozygous mutant embryos at E17.5. The adult cerebral cortex consists of six primary layers, I-VI, from outside (pial surface) to inside (white matter); layer I is the upper most layer adjacent to the pial surface and layer VI is the deepest layer bordering the white matter. Molecular markers, known to label neurons in specific layers (reviewed in (Molyneaux et al., 2007), were used to evaluate the number of neurons in select cortical layers. For example, CTIP2 and TBR1 label neurons in deeper cortical layers (CTIP2 predominantly layer V, with some labeling in layer VI; TBR1for the subplate, layer VI, and some neurons in layer V), while SATB2 and REELIN label neurons in upper cortical layers (SATB2 is for layers II/III and IV/V; REELIN for layer I) (Molyneaux et al., 2007, Colasante et al., 2015). Using antibodies against these four proteins, we compared the lamination of *Snap29* homozygous mutant cortex to that of wild type litter mates. No significant differences were found between genotypes (Figure 8B-E). As embryonic brains have less well-defined layer structures, instead of using layer numbers we used a grid system that divides the embryonic cortical neuroepithelium into eight parts (each with equal distance; grid 1 is closest to the pial surface and grid 8 is closest to the subventricular zone) for quantification (Figure 8B-E). Thus, cerebral abnormalities found in CEDNIK are not modeled in *Snap29* homozygous mutant mice on a mixed genetic background.

However, since we serendipitously observed seizures in a subset of *Snap29* homozygous mutant mice at P10 (N=2/40; Supplemental Video 1), we postulate that these animals may have cerebral malformations but at very reduced penetrance or suffer from neuronal degeneration. Therefore, MRI was used to examine brains of 6-weeks old wildtype (n=3) and *Snap29* homozygous mutant (n= 3) mice (Supplemental Figure 10). However, no significant differences were found, suggesting that severe brain malformations are not responsible for motor defects found in *Snap29* homozygous mutant mice.

**Figure 10.**
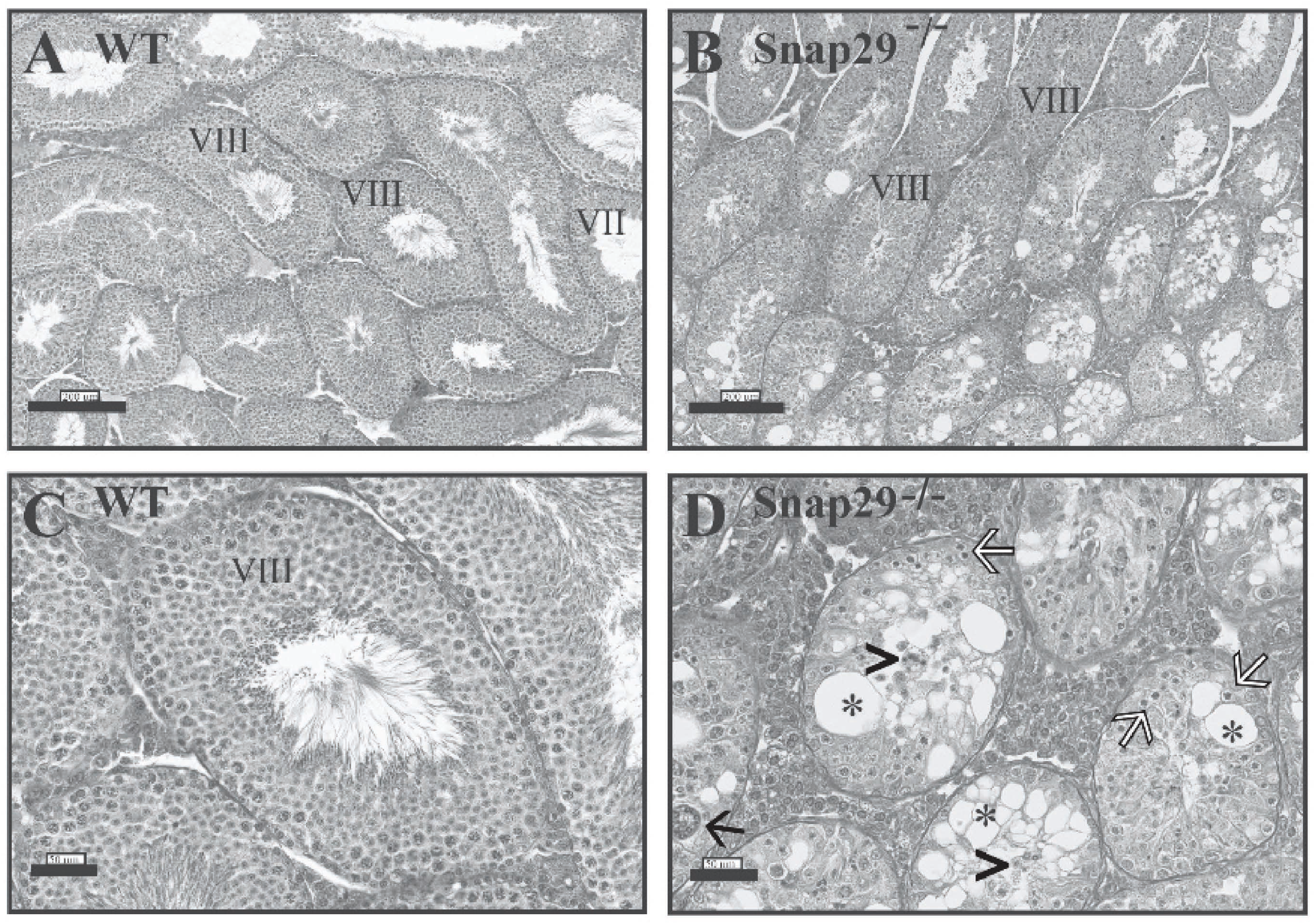
Histological analysis of *Snap29*^-/-^ testis. Testis sections were stained with hematoxylin and eosin to evaluate spermatogenesis in WT (A and C) and *Snap29*^-/-^ (B and D) mice. WT testis displayed normal spermatogenesis (Stages VII and VIII showing elongating spermatids and spermatozoa in the lumen). Seminiferous tubules in *Snap 29*^-/-^ testis had degenerated germ cells (white arrows), loss of immature germ cells accumulated in the lumen (arrow heads), giant multinucleated spermatids (black arrow) and extensive vacuolization (*). The diameter of degenerated seminiferous tubules was reduced in *Snap 29*^-/-^ (D) compared to WT (C) testis.

### Retinal Defects in Snap29 homozygous mutant mice

As a subset of CEDNIK patients also show ophthalmological abnormalities such as optic nerve hypoplasia and atrophy, we used electroretinogram (ERG) to assess retinal function in 21 mice, including 5 wild type, 8 *Snap29* heterozygous and 8 *Snap29* homozygous mutant mice (with 5 males and 16 females whose ages ranged between 75-118 days). An attenuation of the b wave in both scotopic and photopic ERG was observed in 3 out of 8 mice (37.5%) of the homozygous mutant group. The average b wave amplitude of the three mice with abnormal ERG is 319.89±111.44μV in scotopic and 20.95±32μV in photopic, which are significantly lower than those of wild type mice (628.97±113.45μV in scotopic and 107.82±35.6μV in photopic). However, the ERG results of the remaining homozygous and heterozygous mice closely matched the results obtained from the wild type group (Table 5). Figure 9A-C shows representative scotopic and photopic ERG waveforms of wild type, heterozygous and mutants (with low b amplitude). In addition, H&E revealed a significant thinning of the outer nuclear (ONL) and inner nuclear (INL) layers of the retina of *Snap29* homozygous mutant mice compared to the wild type group (40.90 ± 1.69μm vs. 46.15μm, p<0.05 in ONL and 30.87 ± 1.26μm vs. 34.87μm, p<0.05 in INL) (Figure 9D-E). Thus, the *Snap29* homozygous mutant mice model ophthalmological abnormalities found in CEDNIK although with reduced penetrance.

**Table 5:** ERG b wave amplitude average

|  | +/+ (n=5) | +/- (n=8) | -/- (n=3 with abnormal ERG) | -/- (n=5) |
| --- | --- | --- | --- | --- |
| Scotopic | 628.97±113.45μV | 654.31±42.22μV | **319.89±111.44μV** | 672.23±154.04μV |
| Photopic | 107.82±35.6μV | 111.77±12.25μV | **20.95±32μV** | 88.3±12.68μV |

### Snap29 homozygous mutant males are infertile

During our studies, we mated *Snap29* homozygous mutant females and males to generate mutants, however, no live births were found from these mating. To determine if *Snap29* female or male were infertile, we set up mating of *Snap29* homozygous mutant males and females with wild type or heterozygous males and females for 3.9 months. Live births were found in all cases except when mating pairs consisted of a *Snap29* homozygous mutant male mouse (n=4) regardless of the genotype of the female: wild type (n=3) or homozygous mutant females (3), indicating that *Snap29* homozygous mutant males are infertile (Table 6). Importantly, mutant male mice were able to mount and generate vaginal plug in females, suggesting that the weaker grip and or lack of coordination is not the cause of this infertility.

**Table 6:** Fertility of Snap 29 mutant mice

| **mating #** | **male genotype** | **female genotype** | **duration of mating** | **# litters** |
| --- | --- | --- | --- | --- |
| **1** | **wt** | **wt** | **5 months** | **8** |
| **2** | **wt** | **wt** | **5 months** | **8** |
| **3** | **wt** | **wt** | **2 months** | **4** |
| **4** | **wt** | **mut** | **3 months** | **6** |
| **5** | **wt** | **mut** | **4 months** | **6** |
| **6** | **htz** | **htz** | **5 months** | **13** |
| **7** | **htz** | **htz** | **5 months** | **13** |
| **8** | **htz** | **htz** | **3 months** | **9** |
| **9** | **htz** | **htz** | **3 months** | **9** |
| **10** | **htz** | **wt** | **3 months** | **0** |
| **11** | **htz** | **mut** | **3 months** | **0** |
| **12** | **hmz** | **wt** | **4 months** | **0** |
| **13** | **hmz** | **mut** | **4 months** | **0** |
| **14** | **hmz** | **wt** | **4 months** | **0** |
| **15** | **hmz** | **mut** | **4 months** | **0** |
| **16** | **hmz** | **wt** | **3 months** | **0** |
| **17** | **hmz** | **mut** | **3 months** | **0** |
| **18** | **hmz** | **wt** | **3 months** | **0** |
| **19** | **hmz** | **mut** | **3 months** | **0** |

In light gray: mating where no litter were observed

Analysis of the reproductive organs of these males revealed that the testis/body ratio in *Snap29* homozygous mutant mice were significantly less than that of *Snap29* heterozygous or wild type males (0.29×10^-2^ vs 0.50×10^-2^ and 0.54×10^-2^, respectively; p≤0.05, Kruskal-Wallis ANOVA). We did not observe differences in the weight of epididymis, seminal vesicles, coagulating glands and prostate among the three genotypes (Table 7). In addition, a subset of testis of *Snap29* homozygous mutant mice revealed abnormal spermatogenesis with abnormal seminiferous tubules with degenerated germ cells, extensive vacuolization, loss of immature germ cells, and giant multinucleated cells (Figure 10). Furthermore, the percentage of abnormal seminiferous tubules were higher in testis of *Snap29* homozygous mutant males when compared to heterozygous or wild type mice (10.31±3.67, 2.23±1.06 and 0.15±0.10, respectively; p≤0.05, Kruskal-Wallis ANOVA). In addition, the diameter of degenerated seminiferous tubules was reduced in *Snap29* homozygous mutant (Figure 10D) when compared to wild type (Figure 10C), and few seminiferous tubules of testis of *Snap29* homozygous mutant mice had spermatozoa in their lumen. Thus, SNAP29 is required for male fertility and specifically for spermatogenesis.

**Table 7:**
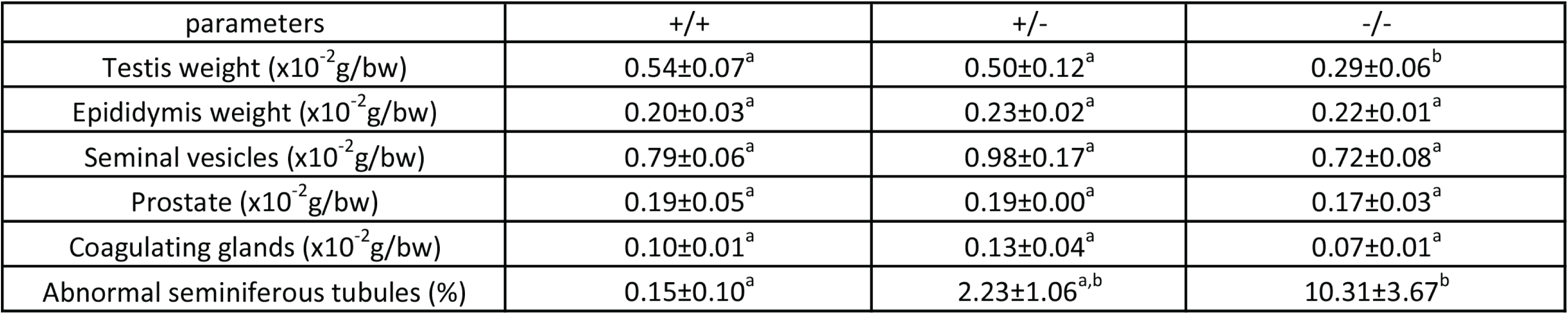
Scored parameters for the organs of male Snap29 mice. Different letters denote significant differences for comparison of each organ among the three groups (Kruskal-Wallis ANOVA, p≤0.05, n=6-8)

| parameters | +/+ | +/- | -/- |
| --- | --- | --- | --- |
| Testis weight ($\times 10^{-2}$ g/bw) | $0.54 \pm 0.07^a$ | $0.50 \pm 0.12^a$ | $0.29 \pm 0.06^b$ |
| Epididymis weight ($\times 10^{-2}$ g/bw) | $0.20 \pm 0.03^a$ | $0.23 \pm 0.02^a$ | $0.22 \pm 0.01^a$ |
| Seminal vesicles ($\times 10^{-2}$ g/bw) | $0.79 \pm 0.06^a$ | $0.98 \pm 0.17^a$ | $0.72 \pm 0.08^a$ |
| Prostate ($\times 10^{-2}$ g/bw) | $0.19 \pm 0.05^a$ | $0.19 \pm 0.00^a$ | $0.17 \pm 0.03^a$ |
| Coagulating glands ($\times 10^{-2}$ g/bw) | $0.10 \pm 0.01^a$ | $0.13 \pm 0.04^a$ | $0.07 \pm 0.01^a$ |
| Abnormal seminiferous tubules (%) | $0.15 \pm 0.10^a$ | $2.23 \pm 1.06^{a,b}$ | $10.31 \pm 3.67^b$ |

## Discussion

Herein we report that abnormalities associated with loss of function mutations of *SNAP29* in patients with CEDNIK or 22q11.2DS are phenocopied in a *Snap29* mutant mouse line on a mixed genetic background. We showed that *Snap29* was ubiquitously expressed during mouse development, which is surprising considering the specific defects associated with mutations in human. Furthermore, our comprehensive analysis of *Snap29* homozygous mutant mice revealed that many more abnormalities associated with CEDNIK syndrome such as motor defects, skin and retinal defects can be modeled in this organism. Our current analysis also revealed a previously unknown role for *Snap29* in male fertility. Since neonatal death of *Snap29* homozygous mutant pups on the inbred C57 genetic background are due to a compromised skin barrier, we postulate that modifiers on the mixed genetic background rescued this defect. We propose that lethality of this mutant mouse model on the mixed genetic background similar to those of human CEDNIK patients is independent of the role for SNAP29 during skin epidermal differentiation.

### Mixed genetic background facilitates better model of CEDNIK

Mouse models are gold-star experimental models of human genetic conditions, and most abnormalities associated with a given human syndrome can be modeled when the orthologous gene is mutated in mice (Jerome and Papaioannou, 2001, Rajagopal et al., 2007). However, it is also clear that genetic background can influence expressivity and penetrance of these abnormalities (Doetschman, 2009). Thus, it is recommended that genetically engineered mouse models be analyzed on mixed genetic background, so as to identify any major roles played by differences in genetic backgrounds (Doetschman, 2009). Since only skin abnormalities were reported in mouse models with loss of function mutation in *Snap29* on an inbred C57Bl/6 mouse, we generated and analyzed *Snap29* homozygous mutant mice on a mixed CD1; FvB genetic background to identify the influence, if any, of genetic background on the penetrance and expressivity of abnormalities associated with CEDNIK syndrome. On the mixed genetic background, the majority of *Snap29* homozygous mutant mice survived the perinatal period and could be studied for other abnormalities associated with CEDNIK syndrome. Our data indicates that genetic background does contribute to expressivity and penetrance of mutations associated with CEDNIK in the mouse model and also revealed unanticipated requirements for *Snap29* in the eye and the reproductive tract.

### Mice with loss of function mutation of Snap29 models a large number of abnormalities associated with CEDNIK syndrome

Unlike *Snap29* homozygous mutant mice on the inbred C57Bl/6 which all develops severe barrier defects associated with abnormal epidermis differentiation, only 82% of *Snap29* homozygous mutants on a mixed genetic background develop skin scaling. Furthermore, barrier defect was not found in any of these mice, suggesting that modifiers on the B6 genetic background predispose mice with epidermal defects to disrupted skin barriers and neonatal death. Furthermore, as CEDNIK patients appear to die primarily due to pulmonary associated diseases (Hsu et al., 2017) despite skin abnormalities, further elucidation of the cause of death of the *Snap29* mutant mouse model on a mixed genetic *Snap29* might help pinpoint the mechanism leading to lethality of CEDNIK patients. Although we cannot formerly rule out that pups which died within the first 3 days of life developed compromised skin barriers at the time of death, the fact that a small number of homozygous mutant pups died without any obvious skin defects suggest that death of *Snap29* homozygous mutant mice on the genetic background may be due to neurological defects, similar to what was found in the zebrafish model (Mastrodonato et al., 2019).

We showed psychomotor delay, motor abnormalities and hypotonia in *Snap29* homozygous mutant mice. However, this was not associated with defects during neurogenesis or associated with gross malformations in adults. This was surprising since CEDNIK patients show severe neurological defects, including polymicrogyria and degeneration of the corpus collosum (Fuchs-Telem et al., 2011, Hsu et al., 2017, Sprecher et al., 2005). Thus, we postulate that motor defects in *Snap29* homozygous mutant mice may be due to motor neuron degeneration, motor neuron projection, or synaptic transmission. A role for SNAP29 in motor neuron branching is supported by work in zebrafish where altered motor neuronal projections was found in ENU mutants and morphants (Mastrodonato et al., 2019). We also uncovered attenuation of the ERG signal and thinning of the retinal structures of *Snap29* homozygous mutant mice. Our findings suggest that retinal defects found in CEDNIK patients may reflect a primary role for SNAP29 in eye development.

Unexpectedly, we found that all *Snap29* homozygous mutant males were infertile. However, infertility could not be explained simply by severe abnormalities found in seminiferous tubules, since only 10% of tubules were abnormal. SNAP29 plays a crucial role in SNARE complex assembly during intracellular membrane fusion events, and is implicated in synaptic transmission. It is postulated that SNAP29 inhibits SNARE disassembly and mediates neurotransmitter release (Su et al., 2001, Pan et al., 2005). During vesicular transport, endocytosis and exocytosis, SNAREs self-assemble into stable four-helix bundles between vesicular membranes and target membranes to mediate fusion of vesicle membranes and to the target membrane (Sprecher et al., 2005).Trans-SNARE complex formation is also important for membrane fusion and completion of the acrosome reaction, a necessary event for fertilization (Sharif et al., 2017). Hence, we hypothesize that SNAP29 is required for spermatogenesis and molecular mechanisms associated with the acquisition of fertilizing ability by the spermatozoon. Although these results are interesting, the role of SNAP29 in male fertility was not the scope of the present study and further investigation is required to extensively elucidate the participation of SNAP29 in spermatogenesis and sperm function.

### Variable expressivity and incomplete penetrance explain contribution of SNAP29 mutation to 22q11.2DS

Interestingly, we observed incomplete penetrance of the *Snap29* mutation on the mix genetic background. We previously showed that deletion of the 22q11.2 region can uncover pathogenic mutations in *SNAP29*, leading to CEDNIK phenotype in a subset of 22q11.2DS patients (McDonald-McGinn et al., 2013). Intriguingly two patients with mutations in *SNAP29* and hemizygous for 22q11.2 were atypical and did not have any skin abnormalities, a hallmark of CEDNIK, while only one patient had neurological abnormalities (McDonald-McGinn et al., 2013). Thus, we proposed that mutations of *SNAP29* may also show incomplete penetrance. This is further supported by the recent report by Sun et al (Sun et al., 2016) of an ADNFLE patient carrying a truncating mutation in *SNAP29.* However, since abnormalities and pathologies have not been reported in parents of CEDNIK patients whom are obligated carriers of *SNAP29* mutations, mutation of *SNAP29* shows incomplete penetrance. In addition, our study revealed that mutation of *Snap29* shows incomplete penetrance in mouse. Therefore, we propose that hemizygosity for *SNAP29* in the majority of patients with 22q11.2DS contributes to a subset of abnormalities found in these patients. More specifically our studies suggest that SNAP29 may play a role in eye and motor defects found in 22q11.2DS. As 90% of 22q11.2DS patients exhibit motor delays (Bassett et al., 2011), we propose that our mouse model can be used to investigate the etiology of motor defects in 22q11.2DS and strongly supports *SNAP29* as a candidate gene for these deficits.

Studies using fibroblasts from CEDNIK patients, the previously published mutant mouse model, *C. elegans*, Drosophila, and zebrafish all suggest that SNAP29 is important during epithelial morphogenesis (Morelli et al., 2014, Morelli et al., 2016, Sprecher et al., 2005, Mastrodonato et al., 2019). More specifically, mutations in SNAP29 result in impaired autophagy, abnormal cell division, and apoptosis (Morelli et al., 2014, Morelli et al., 2016, Sprecher et al., 2005, Mastrodonato et al., 2019). It is possible that some or all of these cellular events are disrupted in *Snap29* mutants. Future work will focus on identifying the specific cellular pathways disrupted in *Snap29* mutant cells.

## Acknowledgement

We would like to thank Dr Mitra Cowan, Associate Director of the transgenic core facility at McGill University, Goodman Cancer Center, for the micro-injection experiments. This work was supported by a grant from the Canadian Institutes of Health Research (MOP#142452). We would like to thank Dr. Sebire for help in brain collection and Dr. Braverman for use of the Rotarod. This work would not be possible without help by Mathieu Simard of the McGill Small Animal Imaging Lab. We thank Drs. McDonald-McGinn and Majewski for support and Dr. Beauchamp for reading of the manuscript. LJM is a member of the Research Centre of the McGill University Health Centre which is supported in part by FQRS.

---

**Supplemental Figure 1.**
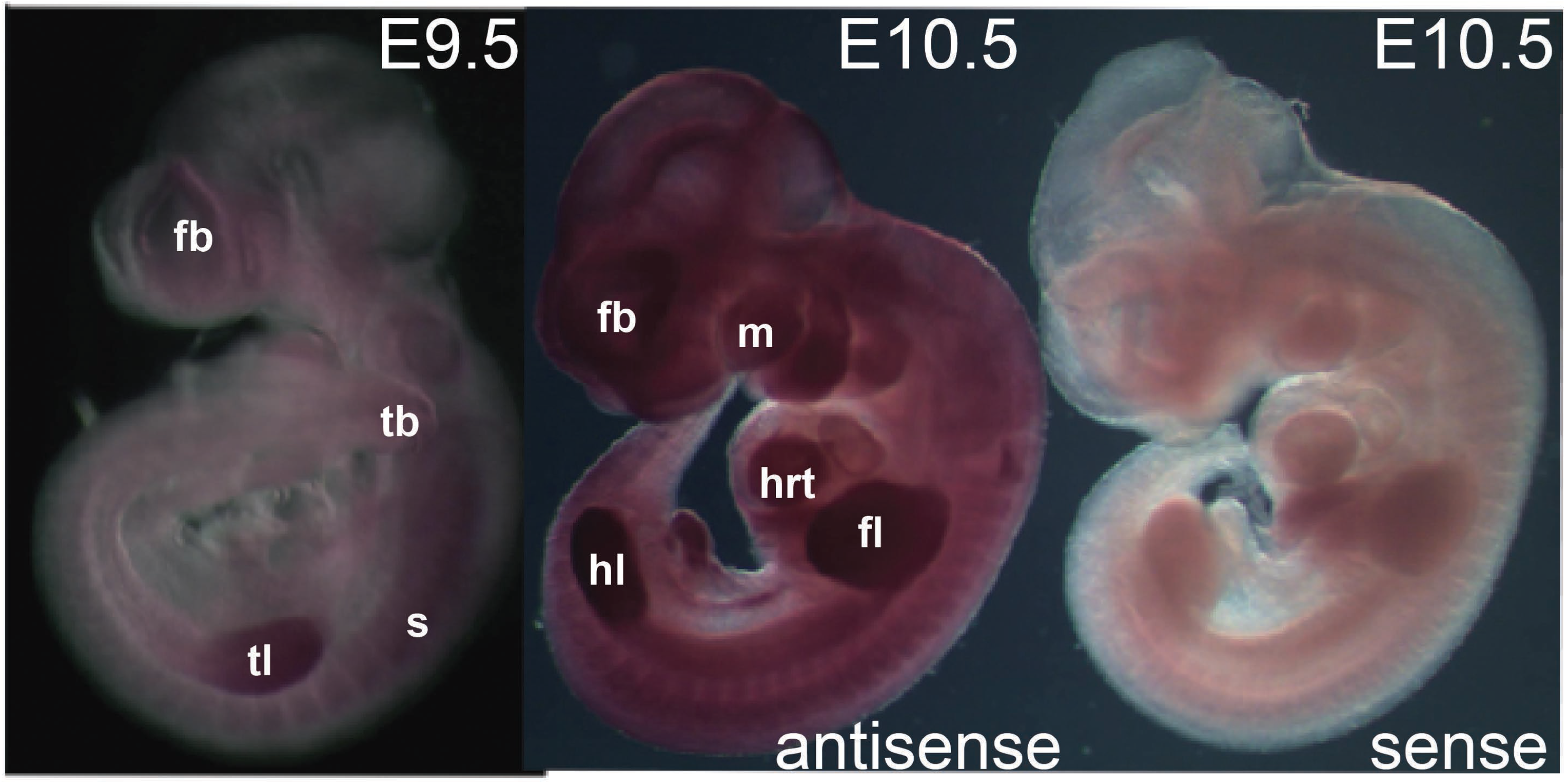
*Snap29* mRNA expression during embryogenesis. *Snap29* mRNA is expressed throughout the embryo at E9.5 including somites and limb buds. By E10.5, expression is detectable everywhere but is more prominent in head tissues and limb buds. fl: fore limb bud, hl: hind limb bud, hrt: heart, s: somite, tb: tailbud, m: maxillary prominence, fb:forebrain

**Supplemental Figure 2.**
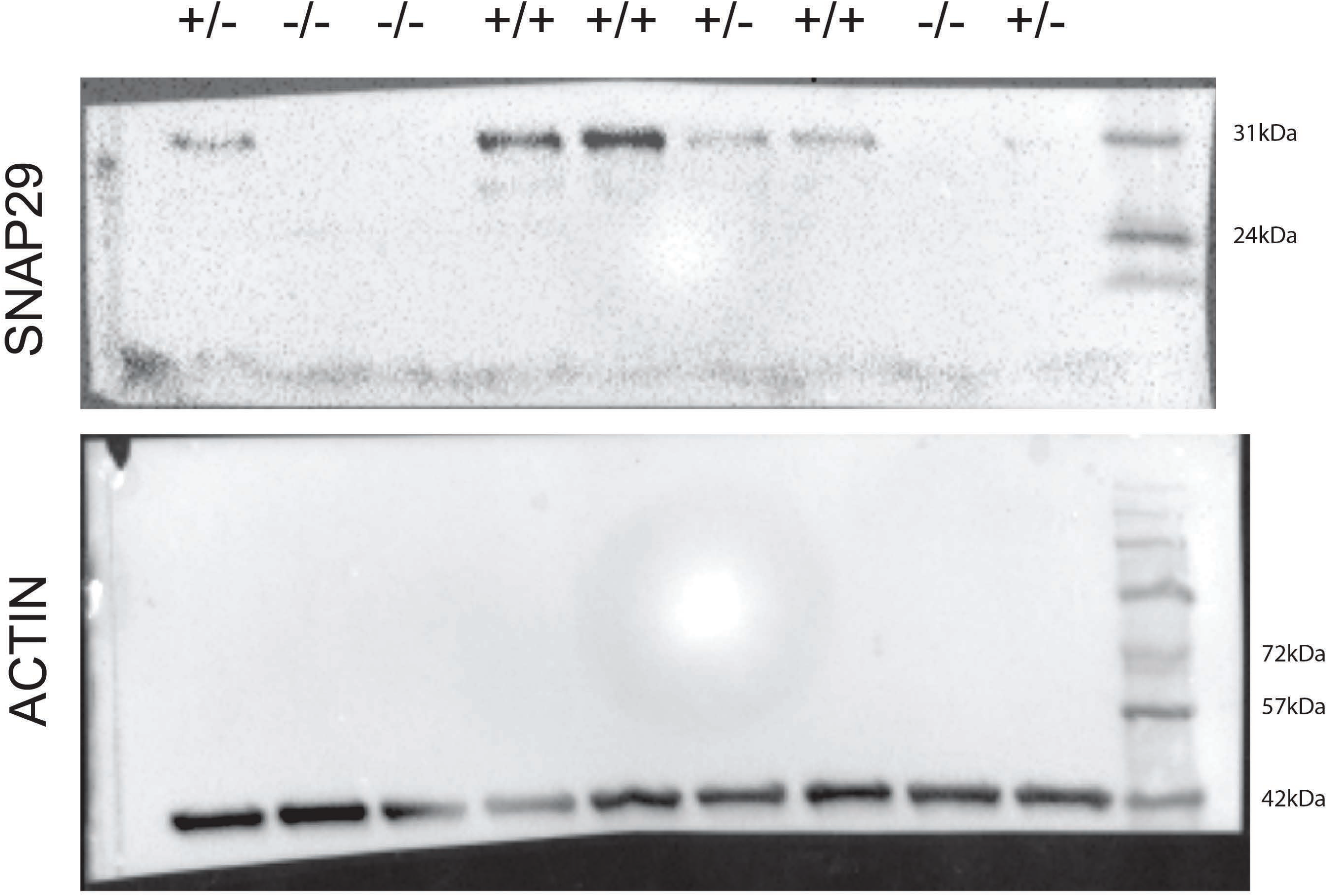
SNAP29 levels in mice carrying the Snap29 allele. Complete western blot shown in figure 1C. Wild-type is denoted by +/+, heterozygotes +/- and homozygous mutants by -/-.

**Supplemental Figure 3.**
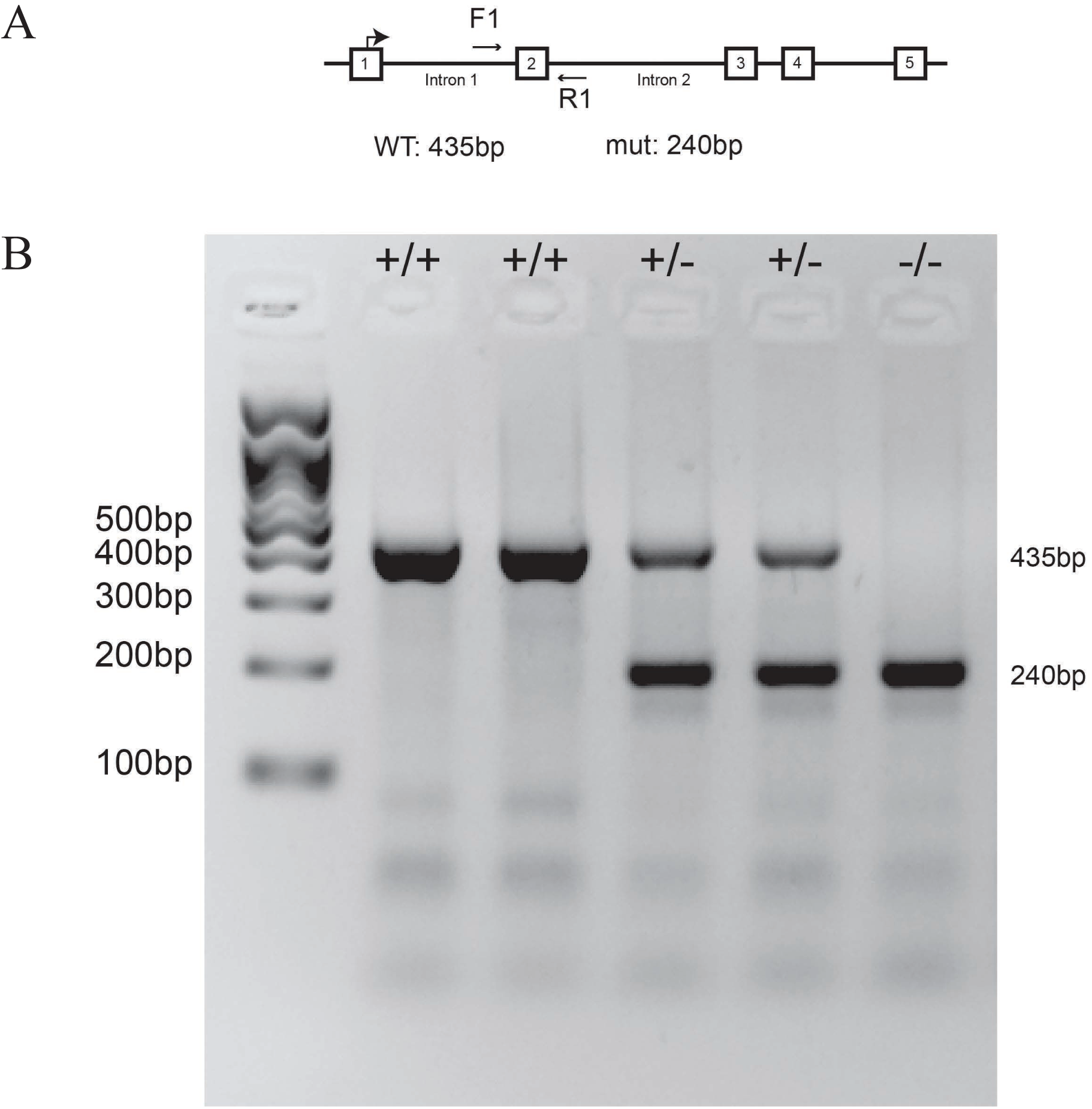
Genotyping of the *Snap29* deletions. A. Schematic representation of the *Snap29* locus with the primer (F1, R1) used for genotyping. B. Example of PCR genotyping for the bigger deletion is depicted here. The WT 435 bp and the mutant 240 bp are detected in heterozygotes.

**Supplemental Figure 4.**
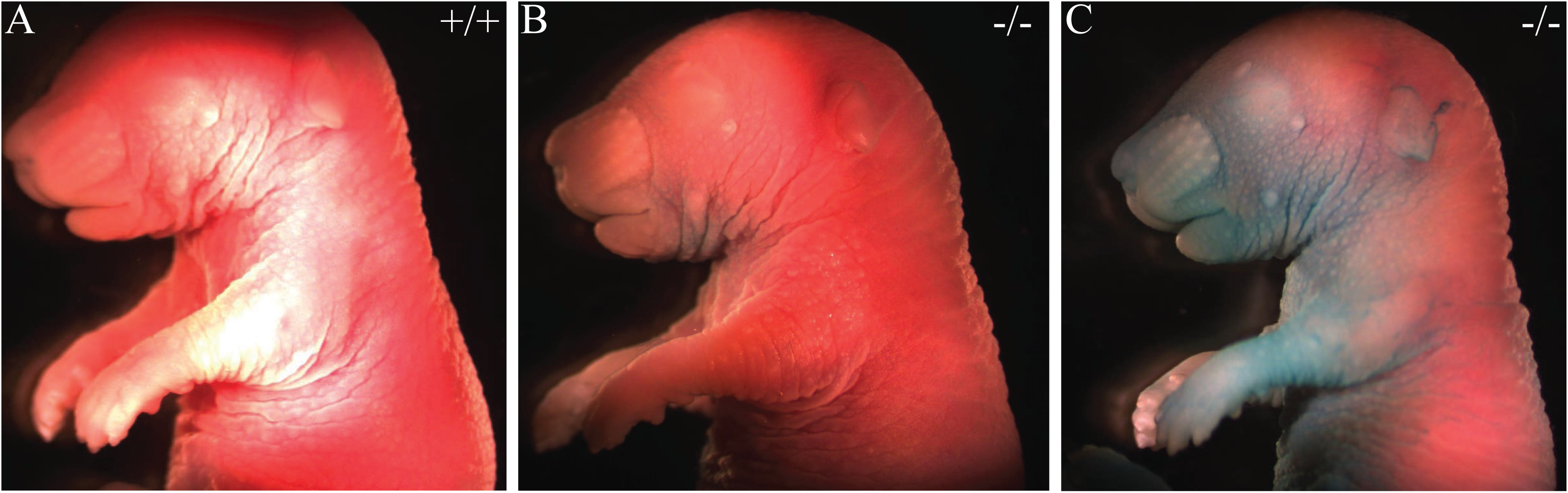
*Snap29* mutant mice show delayed skin barrier formation. X-GAL permeability assay at E17.5. Wild type embryos (A) and some *Snap29*^-/-^ (B) embryos have no coloration suggesting that their skin was not permeable to X-GAL. However, some *Snap29*^-/-^ E17.5 embryos (C) showed X-GAL staining on the ventral body wall, suggestive of a delay in skin barrier formation.

**Supplemental Figure 5.**
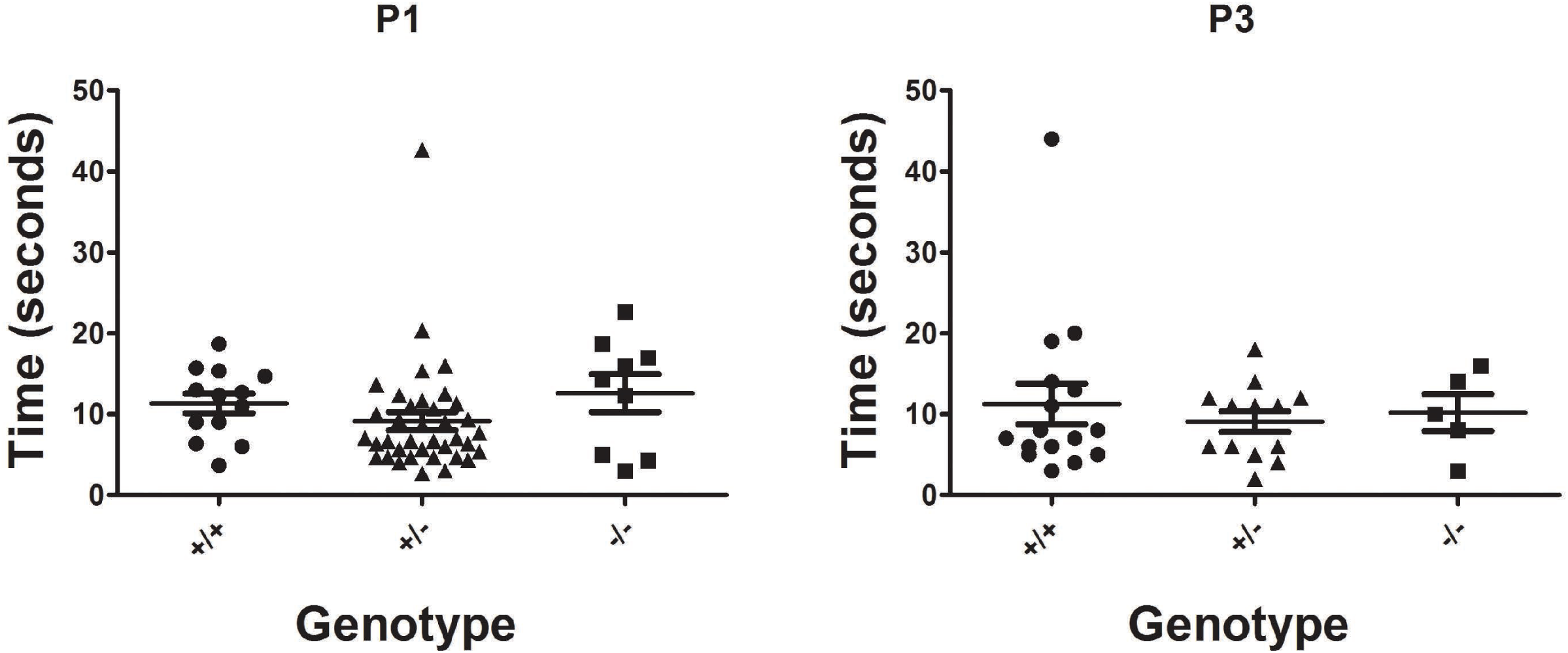
*Snap29* mutant pups turn slightly more slowly than their sibling. The time it took to turn back on their paws when placed on their back in P1 pups (A) and P3 pups (B). The P1 pups turn slightly slower than their sibling although this difference is not significant. By P3 stage, pups are turning at the same rate independently of genotype.

**Supplemental Figure 6.**
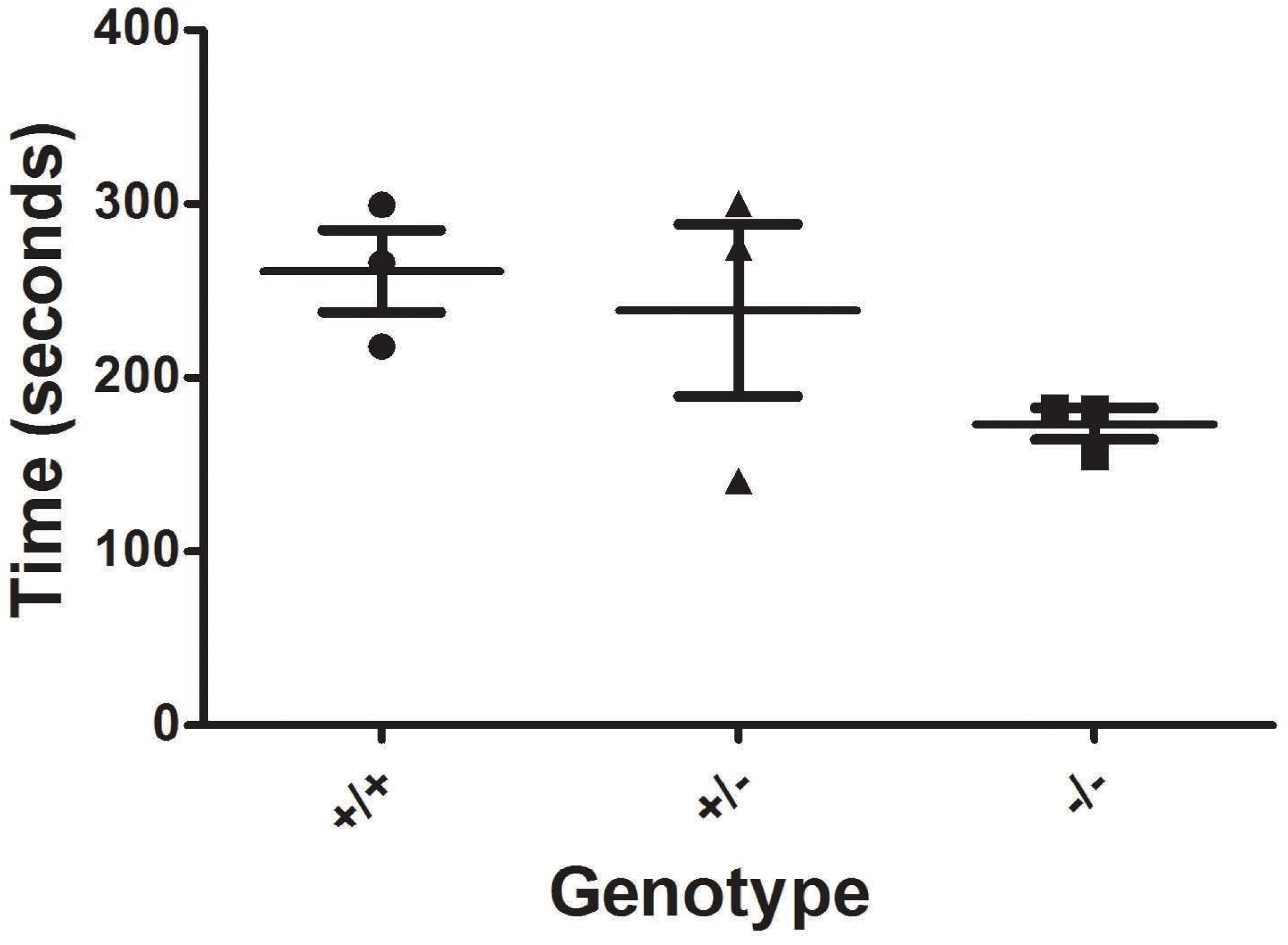
Snap29 mutant animal show a reduced latency to fall. Latency to fall as measured by rotarod assay is more pronounced in *Snap29*^-/-^ male mice, although this result is not significant.

**Supplemental Figure 7.**
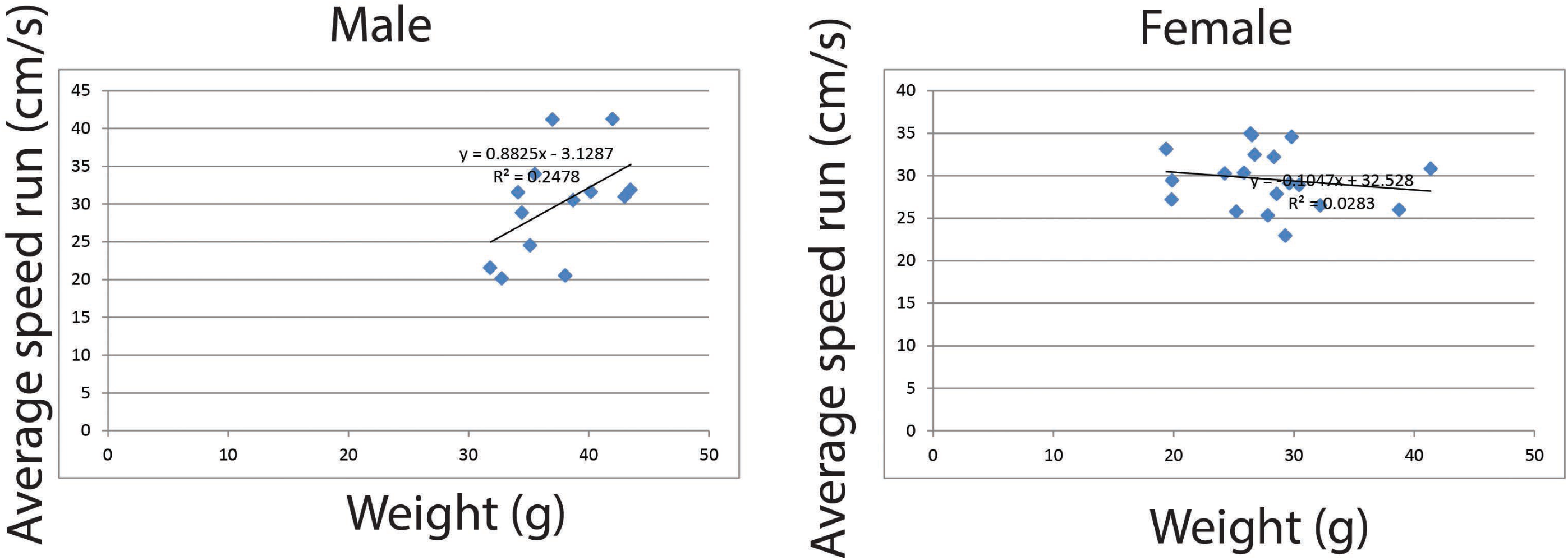
Weight and average run speed are not correlated. A. Plot showing that weight and the speed of male mice are not correlated (R2:0.2478). B. Similarly, the weight and the speed of female mice are not correlated (R2: 0.0283).

**Supplemental Figure 7.**
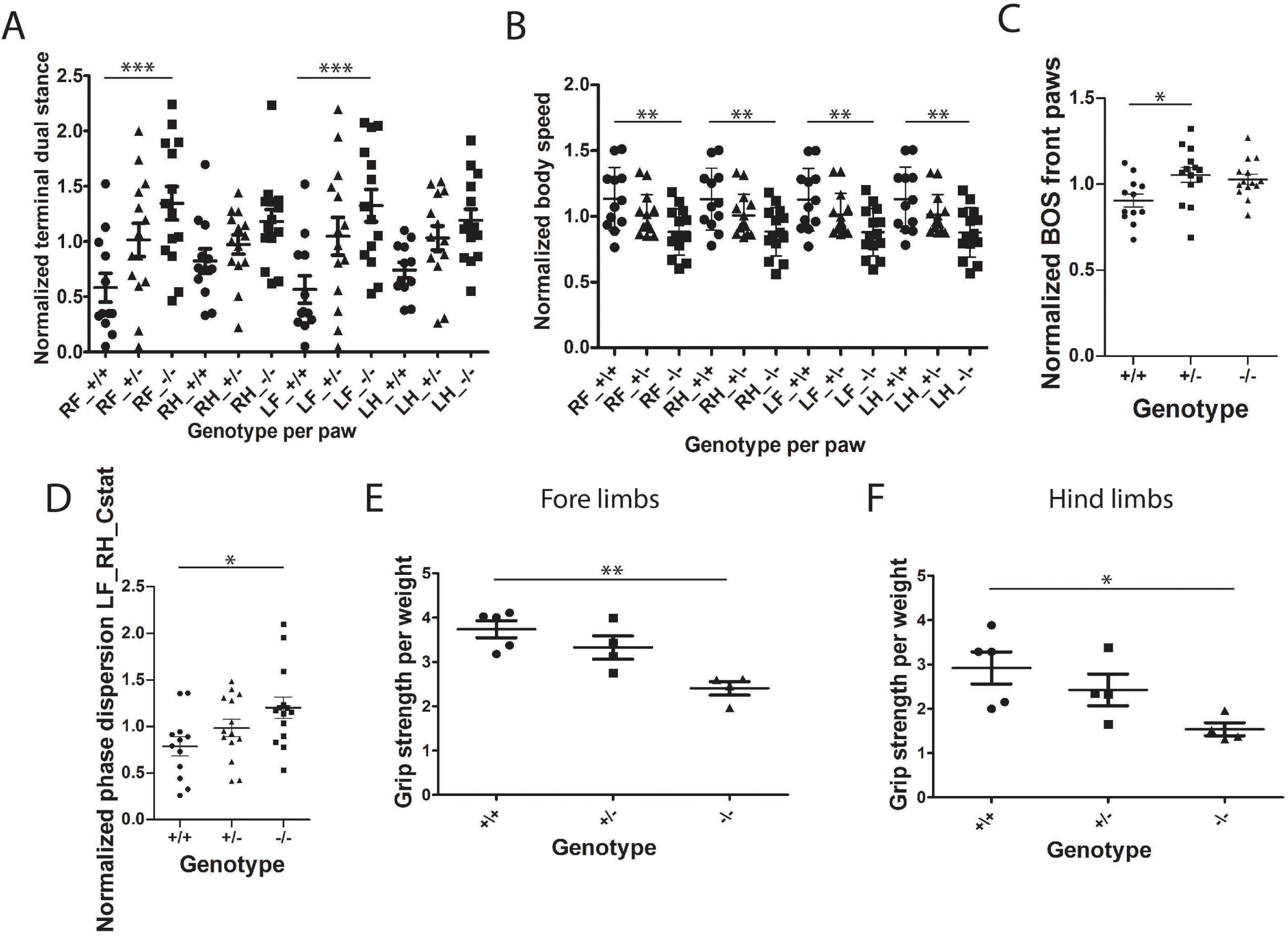
Catwalk gait parameters that were affected in *Snap29*^-/-^ females. Catwalk assay was used to monitor gait parameters in *Snap29*^*+/+*^, *Snap29*^*+/-*^and *Snap29*^-/-^ females. Examples of parameters showing differences in female *Snap29*^-/-^ mutants. The temporal parameter terminal dual stance was significantly elevated in both front paws (A). The kinetic parameter body speed was decrease in all paws of mutant animals (B). The interlimb coordination parameter BOS front paw was significantly increased in *Snap29*^*+/-*^ females when compared to *Snap29*^*+/+*^ animals (C). The interlimb coordination parameters phase dispersion LF_RH Cstat was elevated in mutant females (D). The grip strength was assessed in 14 weeks old female for fore limbs (E) and hind limbs (F). The reduce grip strength did not recover over time in *Snap29*^-/-^ females. RF: right front paw; RH: right hind paw; LF: left front paw and LH: left hind paw. Statistical significance: *: p<0.05, **: p<0.01 and ***: p<0.001.

**Supplemental Figure 9.**
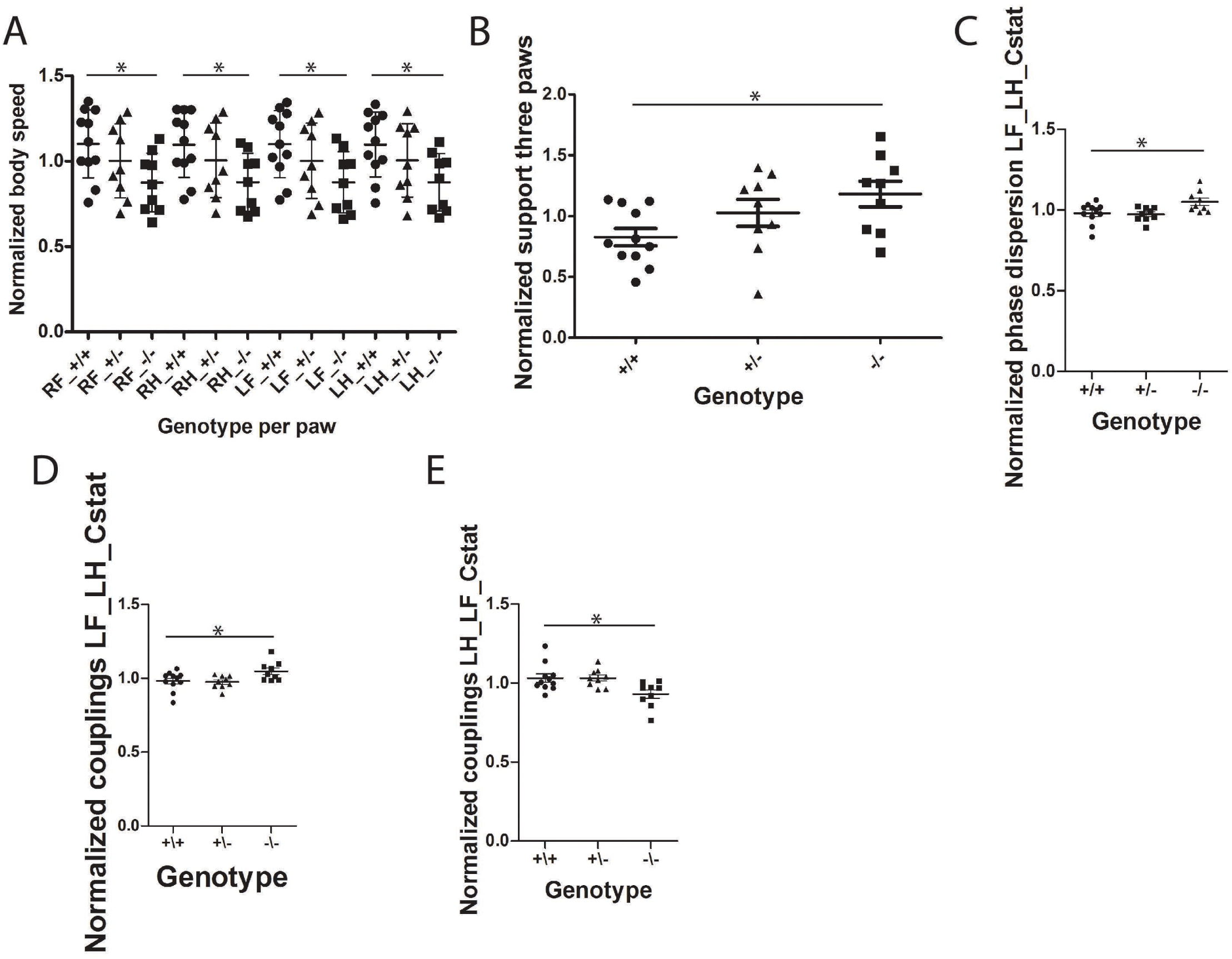
Catwalk gait parameters that were changed in Snap29-/- males. Catwalk assay was used to monitor gait parameters in *Snap29*^*+/+*^, *Snap29*^*+/-*^and *Snap29*^-/-^males. Examples of parameters showing differences in male *Snap29*^-/-^ mutants. The kinetic parameter body speed was decreased for all paws of *Snap29*^-/-^ males (A). The interlimb coordination parameters support on three paws, phase dispersion LF_LH_Cstat and couplings LF_LH_Cstat were significantly elevated in Snap29-/- males (C and D). In contrast, the interlimb coordination parameters couplings LH_LF_Cstat was reduced in *Snap29*^-/-^ males (E). RF: right front paw; RH: right hind paw; LF: left front paw and LH: left hind paw. Statistical significance: *: p<0.05, **: p<0.01 and ***: p<0.001.

**Supplemental Figure 10.**
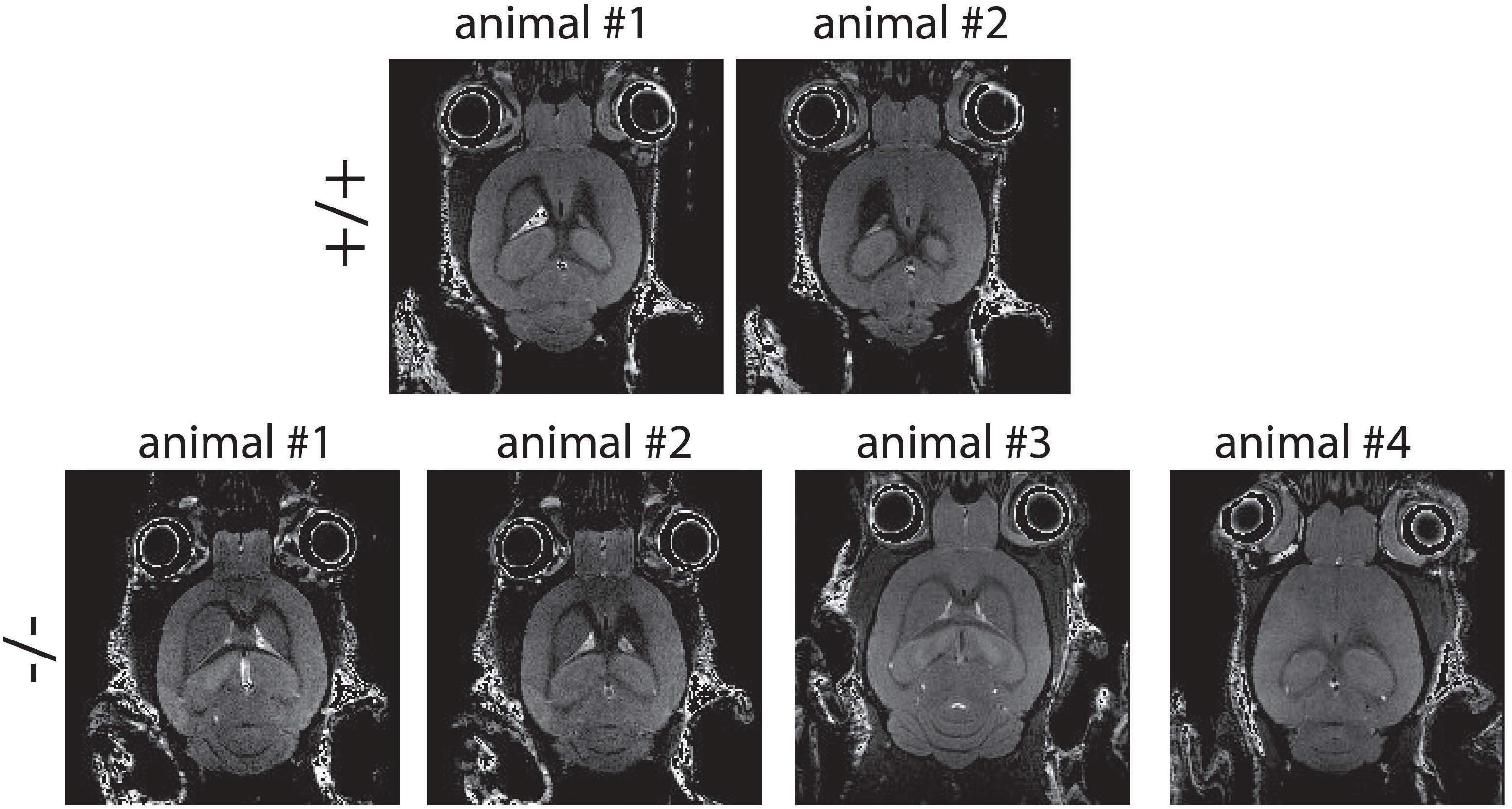
MRI analysis of brain of *Snap29*^*+/+*^ and *Snap29*^-/-^ animals. Representative layers of the head of 11 weeks old animal. Two wild type animals and 4 mutant animals were analyzed. The MRI revealed no differences between these genotypes.

**Supplemental Video S1.**
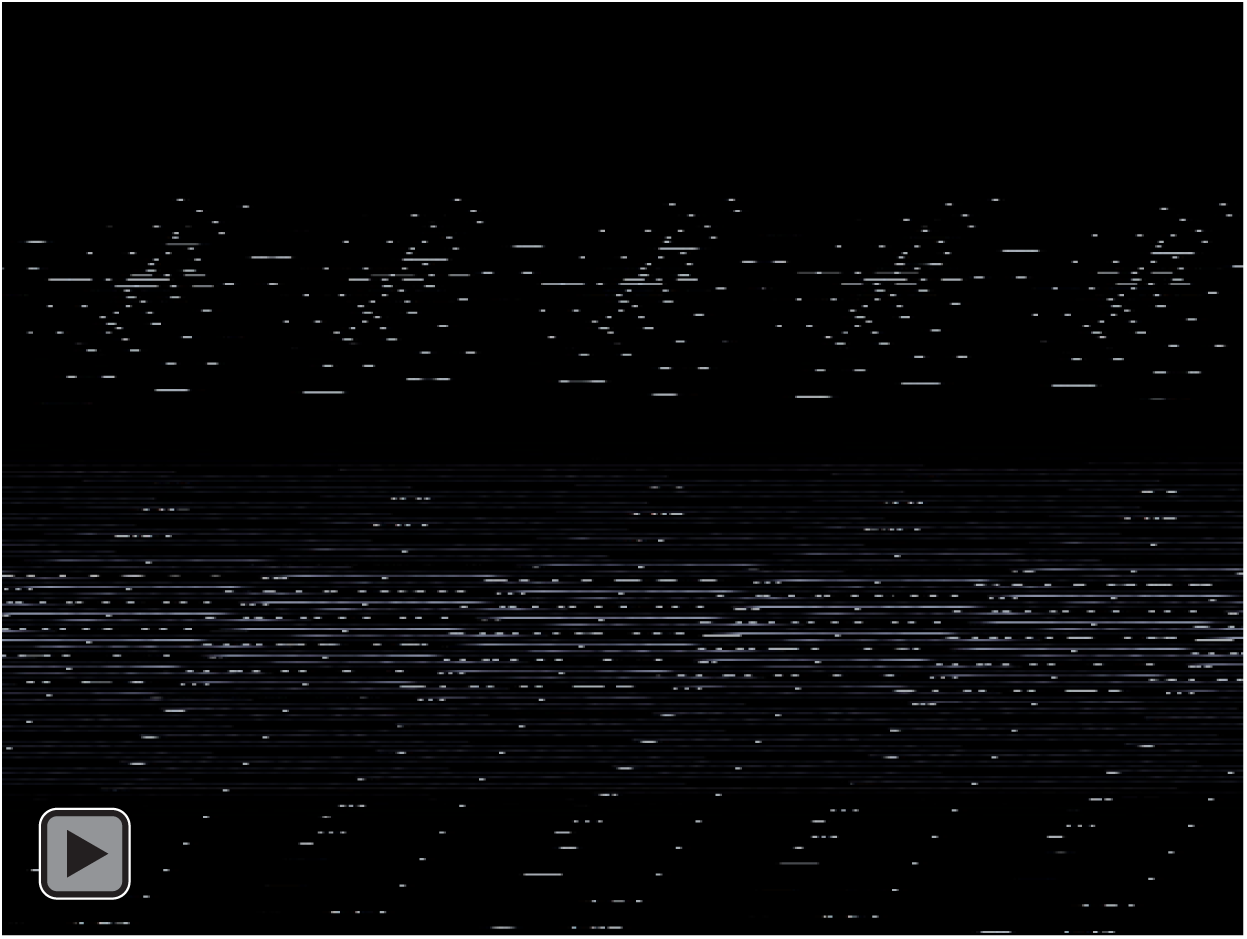
*Snap29*^-/-^ mice exhibiting seizures. Two of the three mice exhibited seizures as measured by episode of intense shaking. These animal turned out to be *Snap29*^-/-^ after genotyping.

